# A genealogy-based approach for revealing ancestry-specific structures in admixed populations

**DOI:** 10.1101/2025.01.10.632475

**Authors:** Ji Tang, Charleston W. K. Chiang

## Abstract

Elucidating ancestry-specific structures in admixed populations is crucial for comprehending population history and mitigating confounding effects in genome-wide association studies. Existing methods for elucidating the ancestry-specific structures generally rely on frequency-based estimates of genetic relationship matrix (GRM) among admixed individuals after masking segments from ancestry components not being targeted for investigation. However, these approaches disregard linkage information between markers, potentially limiting their resolution in revealing structure within an ancestry component. We introduce ancestry-specific expected GRM (as-eGRM), a novel framework for elucidating the relatedness within ancestry components between admixed individuals. The key design of as-eGRM consists of defining ancestry-specific pairwise relatedness between individuals based on genealogical trees encoded in the Ancestral Recombination Graph (ARG) and local ancestry calls and computing the expectation of the ancestry-specific relatedness across the genome. Comprehensive evaluations using both simulated stepping-stone models of population structure and empirical datasets based on three-way admixed Latino cohorts showed that analysis based on as-eGRM robustly outperforms existing methods in revealing the structure in admixed populations with diverse demographic histories. Taken together, as-eGRM has the promise to better reveal the fine-scale structure within an ancestry component of admixed individuals, which can help improve the robustness and interpretation of findings from association studies of disease or complex traits for these understudied populations.

## Introduction

Genetic admixture, the exchange of genetic material of previously relatively isolated populations, results in haplotypes descended from multiple ancestral sources (Korunes & Goldberg 2021; Rius & Darling 2014; Yang & Fu 2018). This phenomenon is pervasive in human populations, exemplified by the genetic admixture experienced by native populations throughout the American continent due to the colonization by Europeans and the subsequent African slave trade (Moreno-Estrada et al. 2013; Conomos et al. 2016). Revealing ancestry-specific structures in admixed populations is crucial for understanding population history and adjusting for population stratification in genome-wide association studies (GWAS). These structures provide insights into migration patterns and genetic diversity, improving our understanding of complex population histories (Moreno-Estrada et al. 2013; Browning et al. 2016). In GWAS, failure to account for population structure can lead to spurious associations or mask genuine genetic effects (Marchini et al. 2004; Martin et al. 2019; Sohail et al. 2019). However, elucidating these structures presents significant challenges due to the intricate genetic composition of admixed individuals, particularly in cases of recent admixture or populations with multiple ancestral sources.

The conventional approach for revealing population structure involves constructing a variance-standardized Genetic Relationship Matrix (GRM) and applying Principal Component Analysis (PCA), at times in conjunction with Uniform Manifold Approximation and Projection (UMAP), to the GRM (Price et al. 2006; Patterson et al. 2006; Novembre et al. 2008; Chiang, Mangul, et al. 2018; Chiang, Marcus, et al. 2018; Diaz-Papkovich et al. 2019; Sakaue et al. 2020; Diaz-Papkovich et al. 2021). In the context of admixed populations, these approaches effectively average over the distribution of ancestral background at a genetic variant and across all loci in the genome, without incorporating ancestry information. Consequently, multiple components of ancestries could mask the finer-scale structure that may be of interest as inter-continental distances tend to dominate and explain the largest amount of variation in the GRM. Therefore, PCA or UMAP applied directly to GRM from admixed individuals tend to reveal structure driven by different proportions of ancestries, even among the lower PCs.

To address this limitation, Moreno-Estrada et al. (Moreno-Estrada et al. 2013) proposed an ancestry-specific PCA method named ASPCA. ASPCA masks genomic components derived from non-target ancestral populations and then compute the subspace spanned by the first k PCs by finding a matrix decomposition that minimizes the reconstruction error (Johnson et al. 2011; Moreno-Estrada et al. 2013). After observing artifactual separation of clusters between reference and admixed individuals when using ASPCA, Browning et al. (Browning et al. 2016) proposed a variant of this ancestry-specific PCA method (we refer to this method as Browning’s Ancestry-Specific Multidimensional scaling, or AS-MDS), which applies MDS to a Euclidean distance matrix based on pairwise allelic differences between individuals after non-target ancestries are similarly masked. Finally, though not yet peer-reviewed, another ancestry-specific PCA method (Missing DNA PCA, mdPCA; https://github.com/AI-sandbox/mdPCA) is also available that constructs a covariance matrix that masks the components with non-target ancestries and then utilized multiple matrix denoising techniques and truncated singular value decomposition on the covariance matrix to compute ancestry-specific PCs. In all these methods linkage information was discarded, and thus these methods are expected to not fully utilize the genomic information for elucidating population structure.

The entire genealogy of the DNA sequence of a sample of individuals can be represented by a series of genealogical trees connected through recombination events, collectively known as the ancestral recombination graph (ARG) (Hudson 1990; Griffiths & Marjoram 1996). With the recent ability to infer or approximate the ARG in thousands of individuals, multiple downstream ARG-based population and statistical genetic applications have been developed to enhance our understanding of the evolutionary history of a population (Lewanski et al. 2024; Brandt et al. 2024; Nielsen et al. 2025). We previously developed an ARG-based framework, called eGRM, to infer the expected relatedness between pairs of individuals (Fan et al. 2022). eGRM utilizes the same variance-standardized framework as the canonical GRM but sums over the vector of haploid individuals for each branch, weighted by branch lengths. As this approach leverages haplotype information to infer the ARG, it enhances robustness when working with incomplete genetic data and improves over canonical GRM in elucidating the population structure of a sample through PCA and UMAP. However, eGRM does not remove the components with non-target ancestries, limiting its application to detect ancestry-specific structure in admixed populations.

In this study, we propose as-eGRM, a framework that integrates ARGs and local ancestry information to infer the expectation of pairwise genetic relatedness within ancestries in an admixed population. We show that PCA and UMAP applied to as-eGRM can outperform alternative methods such as AS-MDS and mdPCA in revealing ancestry-specific structures in admixed populations. We used simulated data of varying complexity to extensively evaluate the performance of as-eGRM on revealing the finer structure in admixed populations. Finally, we applied as-eGRM to a real-world dataset of admixed Latino populations from the HCHS/SOL dataset and the PAGE-Latin American dataset.

## Material and methods

### Expected pairwise genetic relatedness based on genealogical trees

We first briefly review the definition and construction of the eGRM, which provides the pairwise genetic relatedness with a genealogical tree (Fan et al. 2022). Given a branch *e* on a genealogical tree *t* within an ARG *G*, the eGRM defines the genetic relatedness, *R*^*t*^, between a pair of haplotypes *i* and *j* on a single tree as,

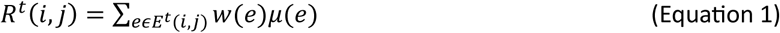

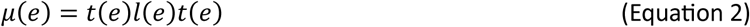

where *E*^*t*^(*i*, *j*) denotes the set of the branches connecting haplotype *i* to haplotype *j* on tree *t* and *w*(*e*) is a weighting function that will be discussed further below. As the number of mutations occurring on each branch *e* of the tree is modeled as a Poisson process, its rate is *μ*(*e*), which is the product of *t*(*e*), *l*(*e*), and *u*(*e*), denoting the length of branch *e* in generations, the number of base pairs that the tree *t* covers, and the mutation rate on this branch, respectively. We use *x*(*e*) to denote the haplotype vector (vector of haploid individuals) associated with branch *e*, that is,

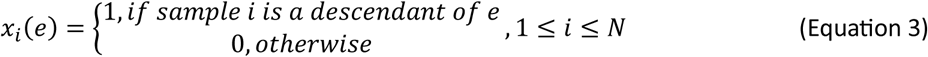

To computationally implement *R*^*t*^, we traverse each branch *e*, compute *w*(*e*)*μ*(*e*) and add *w*(*e*)*μ*(*e*) to the elements in *R*^*t*^ indexed by the descendant samples of branch *e*:

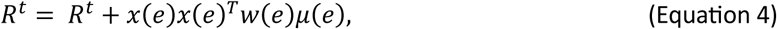

Therefore, across all haplotypes,

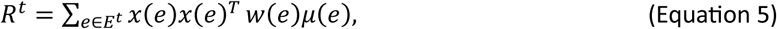

Finally, the relatedness measure is averaged across all trees in the ARG, *G*. With centering, the eGRM is finally defined as:

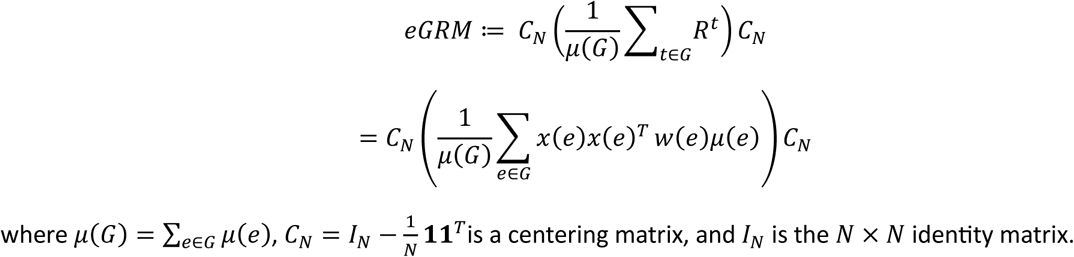

where 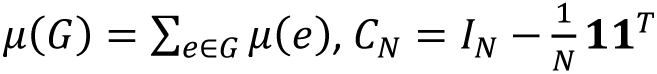 is a centering matrix, and *I*_*N*_ is the *N* × *N* identity matrix.

### Expected pairwise genetic relatedness with ancestry-specific genealogical trees

We define as-eGRM as the eGRM computed on ancestry-specific trees within *G* (**Figure 1A**). By intersecting with the local ancestry information, we prune haplotypes from the tree *t* that are not from the ancestry of interest and re-define the tree while setting those haplotypes as missing. In other words,

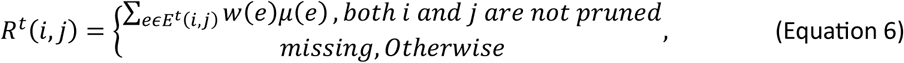

**Figure 1.**
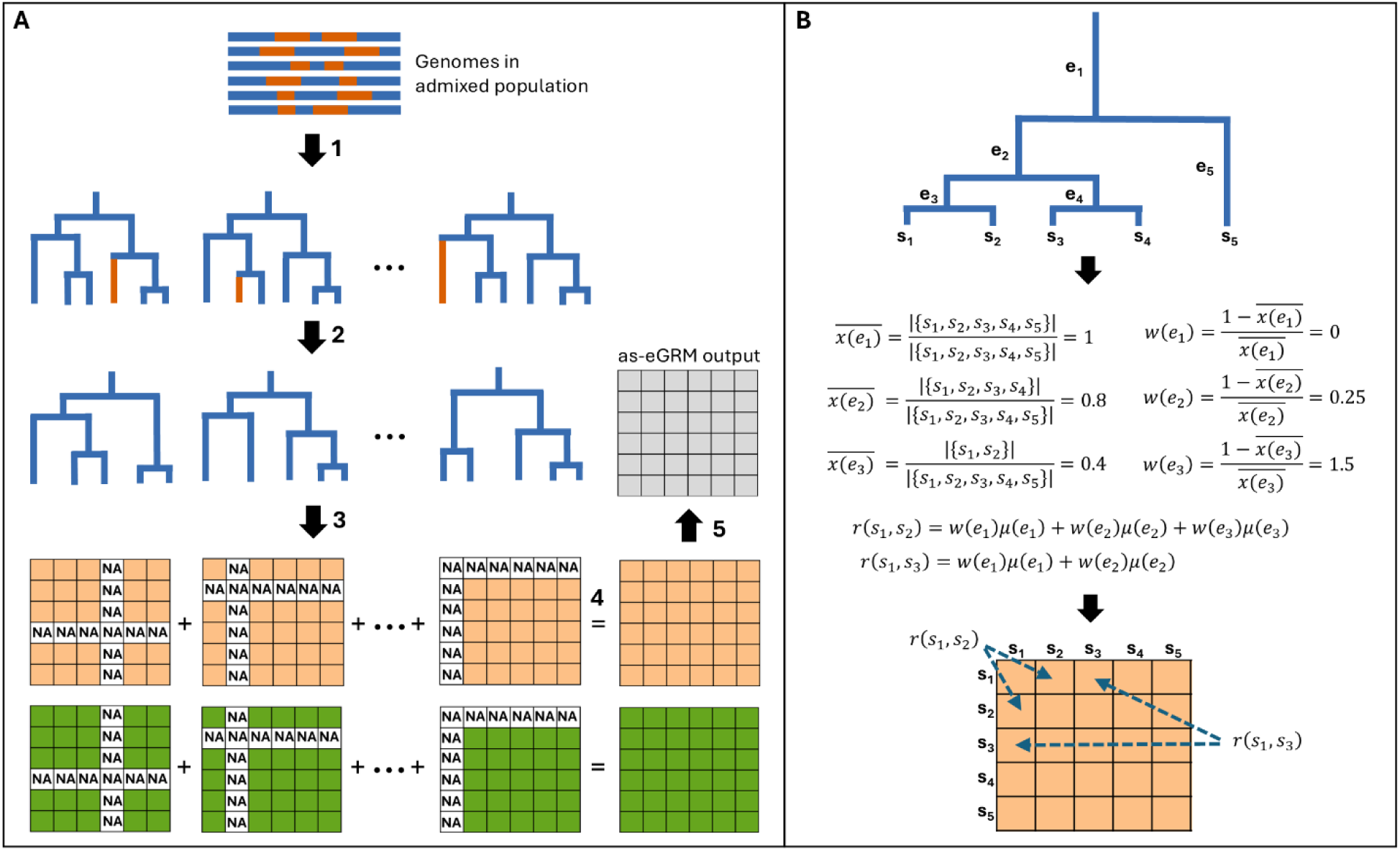
Design of as-eGRM. **(A)** A visual schematic of the implementation of as-eGRM. See the text for detailed description. **(B)** A toy example of the computation of pairwise genetic relatedness. The details are described in the **Method**, Equation (5) under the section (**Expected pairwise genetic relatedness based on genealogical trees**). Here we show a tree with five haplotypes, *s*_1_ to *s*_5_, connected through a tree with five branches, *e*_1_ to *e*_5_. *r*(*s*_*i*_, *s*_*j*_) denote the relatedness between *s*_*i*_ and *s*_*j*_, *μ*(*e*_*i*_) denote the expected number of mutations occurring on branch *e*_*i*_, and 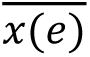 denote the proportion of the descendant samples under branch *e*_*i*_in all the samples. Weights on each branch, *w*(*e*_*i*_), are calculated based on 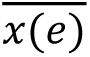, and given the weights and the expected number of mutations we can compute *r*(*s*_*i*_, *s*_*j*_) using all branches that connect the two haplotypes.

We denote the summing matrix across the ARG *G* as *R*^*G*^ = ∑_*t*∈*G*_ *R*^*t*^. As each tree have different number of haplotypes set as missing due to deriving its local ancestry from non-targeted ancestries, instead of dividing the summing matrix by a constant *μ*(*G*), we divide the summing matrix by a *N* × *N* matrix (denoted *D*^*G*^) to account for the differential missing level while taking into account the expected number of mutations occurring on each tree (**Figure 1A**). Each element *D*^*G*^(*i*, *j*) represents the sum of non-missing *μ*(*e*) at position (*i*, *j*) across the *R*^*t*^ (1 ≤ *t* ≤ |*G*|, where |*G*| is the number of trees in *G*):

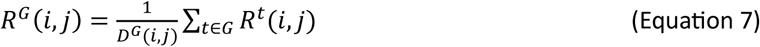

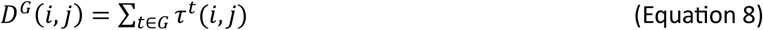

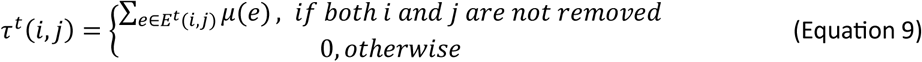

Finally, we center *R*^*G*^ as we would of a regular eGRM:

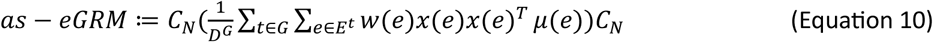

### Choosing the weighting to better reveal recent population structure

The weights on each branch, *w*(*e*), was originally defined in eGRM as 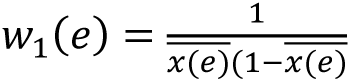, which stem from the canonical GRM term to adjust for the binomial variance of variants across different frequencies (Fan et al. 2022). As *x*(*e*) is the haplotype vector associated with branch *e*, 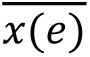 denotes the proportion of the haplotypes under branch *e*. We found that in the context of a genealogical tree, this weight places higher weights on both recent branches (e.g. when 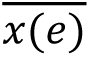 is small, near the leaves of the tree) as well as ancient branches (e.g. when 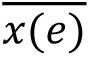 is large, near the root of the tree; **Figure S1A**). Because human population structures are likely established more recently and we tend to be much more interested in the population structure of the recent past (on an evolutionary scale), the original weighting scheme is suboptimal. Indeed, in a simple two-subpopulation two-way admixture model (**Figure S2**), we observe that 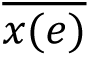 tend to be large for ancient branches, and small for recent branches (**Figure S1C**). Thus, the original weight, *w*_1_(*e*), tend to place higher weights on the more ancient branches particularly when taking into account the longer branches and opportunities for mutations in those branches (**Figure S1D**). We thus experimented with different parametric weighting functions (**Figure S3**) and decided to use weighting function of the form 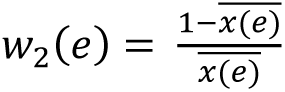 to be effective in up-weighting the more recent past of the genealogical tree (**Figure S1B, S1E**) when computing the expected pairwise relatedness. The software as-eGRM (https://github.com/jitang-github/asegrm) allows users to input different functional forms of the weight.

### Simulation of admixed populations

Three demographic models were used to simulate admixed populations: (1) a two-population split two-way admixture model, (2) a grid-like 3×3 stepping stone model, and (3) a three-way admixed Latino model. For all models we used msprime(version 1.2.0) (Baumdicker et al. 2022) to simulate genetic data with the recombination and mutation rates were set to 1e-8 per generation per base pair, a ploidy of 2 and 500 haplotypes (each spanning 100Mb) per population. In the two-population split, two-way admixture model (**Figure S2**) and the grid-like stepping-stone model (see below), the effective population size was set to 10000 for all populations, with no recent growth. A three-way admixed Latino model (see below) was based on a previously published model that fitted the admixture history model from self-reported Latino Americans from Los Angeles (Fan et al. 2023), but revised to include substructure. A visual representation and detailed parameter specifications are shown in the respective figures and supplementary figures. The commands for simulations are released with the as-eGRM software.

### Quality control of empirical data

We tested as-eGRM and compared it to alternative approaches on empirical data from the HCHS/SOL and PAGE global reference datasets. The HCHS/SOL dataset were obtained from dbGAP (accession numbers phs000880.v1.p1 and phs000810.v1.p1). HCHS/SOL is a large US-based study of 16,415 Hispanic/Latino individuals, among whom 12,803 consented to genetic studies and were successfully genotyped on a genome-wide SNP array (Sorlie et al. 2010). Quality control of genotypes was performed using PLINK (Chang et al. 2015), excluding variants that had a call rate < 99% or P value for Hardy–Weinberg equilibrium < 1.0 × 10^−6^, as well as individuals that has > 2% missingness. For the HCHS/SOL data, we retained only individuals whose four grandparents were self-reported to be from the same country and filtered out relatives by removing one individual in the pairs with kinship (calculated by PLINK) greater than 0.08 (corresponding to second-level relatives or closer). After quality control filtering, we retained 2,036,821 variants and 8,260 individuals for analysis. Among the 8,260 individuals, 1,867 have an estimated Indigenous American ancestry proportion greater than 0.5, which we analyzed to be consistent with the filtering based on ancestry proportion that previous methods used (Browning et al. 2016). Additionally, we also analyzed 1,671 individuals from the Chicago recruitment site across the entire ancestry proportion spectrum, to illustrate the robustness of as-eGRM to missing data for a dataset of similar scale. The PAGE global reference dataset were obtained from dbGAP (accession number phs001033.v1.p1). We extracted a subset of 630 Latin America individuals (from Peru, Venezuela, Mexico, Colombia, Brazil) from the global reference dataset and applied the same quality control filtering to retain 1,399,468 variants for analysis. HCHS/SOL and PAGE-Latin American data were combined with the ancestry reference (see below) and together phased by EAGLE (Loh et al. 2016).

### Inference of Ancestral Recombination Graphs and local ancestry calls

We used Relate (version 1.2.0) (Speidel et al. 2019) to infer ARGs for both simulated and empirical datasets. For simulated data, recombination rate, mutation rate, and effective population size were set to match the simulation parameters. For the HCHS/SOL and PAGE-Latin American data, mutation rate and effective population size were set to the default values as suggested in the user manual, along with the HapMap Phase II genetic map (The International HapMap Consortium 2007) in hg38. For computational scalability when inferring the ARG on empirical datasets, we applied Relate on chunks of 10,000 SNP in parallel. The utility *RelateFileFormats --mode ConvertToTreeSequence* was used to convert Relate’s output to the *tskit* (Kelleher et al. 2018) format.

RFMix (version 2) (Maples et al. 2013) was used to infer local ancestry segments in both simulated and empirical datasets. The ancestral references used in simulation are indicated in each respective simulation model. For running RFMix on the HCHS/SOL and PAGE-Latin American data, we used previously selected individuals based on gnomAD v3.1(Karczewski et al. 2020; Jeon et al. 2023) as the reference. In gnomAD’s nomenclature, we included 671 non-Finnish European (NFE) individuals for European ancestry, 716 African/African-American (AFR) individuals for African-ancestry, and 94 Admixed American (AMR) individuals (7 Colombian, 12 Karitianan, 14 Mayan, 4 Mexican in Los Angeles, 37 Peruvian in Lima, Peru, 12 Pima, and 8 Surui) for Indigenous American ancestry.

### Implementation of previous methods to investigate population structure

We compared PCA + UMAP on the as-eGRM to that based on the canonical GRM, the original eGRM, as well as Browning’s AS-MDS and mdPCA. For PCA on the canonical GRM, we pruned sites with minor allele frequency (MAF) < 0.01 and those in high linkage disequilibrium (LD) using PLINK with the command “--maf 0.01 --indep-pairwise 50 5 0.2”. Then a variance-standardized GRM was computed on the pruned genotypes, followed by eigen-decomposition to derive principal components (PCs). For PCA on the eGRM, eGRM was constructed using the software package from https://github.com/Ephraim-usc/egrm, using the same ARG input as the as-eGRM. Eigen-decomposition was performed on the output of eGRM to compute PCs. For Browning’s AS-MDS and mdPCA, codes were downloaded from https://faculty.washington.edu/sguy/local_ancestry_pipeline/ and https://github.com/AI-sandbox/mdPCA respectively, and executed per instructions from the user manuals. We applied the same MAF and LD pruning as in PCA of the canonical GRM. In particular, mdPCA proposed five different methods (Methods 1-5) for generating ancestry-specific PCs. All five methods were tested, and we found generally the best results were based on Method 1, which were presented in this study. For methods that leverage local ancestry segments (Browning’s AS-MDS, mdPCA, and as-eGRM), used the same local ancestry calls. Unless otherwise noted, UMAP was applied to the top 20 and 50 PCs from simulated and empirical data, respectively, using the Python package *umap* with default parameters (n_neighbors:15, min_dist:0.1, metric: Euclidean).

We visually compare the PCA or PCA+UMAP results based on each method, plotting generally the top two components in a biplot. To quantify the degree of clustering effectiveness, we followed previous study and used the Separation Index (SI), which assess the proportion of nearest neighbors that are in the same population in multi-dimensional space (Fan et al. 2022). Intuitively, for each individual in a cluster of true size *n*, we compute the proportion of the *n* closest neighbor in the multidimensional space that are in the same cluster and average the proportion over all individuals in the dataset. SI is a real number between 0 and 1, indicating how well a multidimensional metric is capturing the true classification. In simulation, the true label is the deme or population membership of each individual. In empirical data, the self-reported country of origin based on grandparental birthplaces in HCHS/SOL or the provided country of origin for PAGE global reference were used as the true label.

## Results

### An overview of the design of as-eGRM

To compute ancestry-specific expectation of genetic relatedness, we first create ancestry-specific trees from inferred genealogical trees. The mathematical formulations are described in detail in the **Methods**. We intersect the inferred genealogical trees with inferred local ancestry segments (in practice inferred from existing methods such as Relate (Speidel et al. 2019) and RFMix (Maples et al. 2013), respectively; **Figure 1A**, step 1). We remove the leaf nodes derived from non-target ancestral populations to generate ancestry-specific trees (**Figure 1A**, step 2). Further, for each of the ancestry-specific trees, we specify two *N* × *N* matrices (named *R*^*t*^ and *τ*^*t*^ respectively; **Figure 1A**, step3; see **Methods**). *R*^*t*^(the orange matrices in **Figure 1A**) scores all pairwise relatedness based on the corresponding tree *t* in the ARG with positions indexed by one or both samples deriving ancestry from the non-target ancestries set to missing values. The pairwise relatedness is computed following the procedure illustrated in **Figure 1B**, which followed the principle of the original eGRM (Fan et al. 2022) that treats mutations as random, and computes the expected relatedness summed across all branches connecting the two haplotypes weighted by the probability of a mutation occurring on the branch (*i.e.* proportional to the branch length; **Methods**). *τ*^*t*^ (the green matrices in **Figure 1A**) records the expected number of mutations on each tree corresponding to the non-missing cells in *R*^*t*^, thereby tracks the differential missingness between pairs of haplotypes due to different proportions of the non-target ancestries being masked across the genome. Across all trees in the ARG, we then take the element-wise sum of the two matrices respectively, producing two matrices *R*^*G*^ and *D*^*G*^(**Figure 1A**, step 4; see **Methods**). Finally, we take the element-wise ratio of the two summed matrices followed by mean-centering to generate the final as-eGRM (**Figure 1A**, step 5).

When computing the expectation of relatedness per branch, the original formuation from the eGRM included a weight based on the inverse of the binomial variance (see **Methods**). This stemmed from the practice in the canonical GRM in which the contribution from each variant is normalized to adjust for the binomial variance of variants across different allele frequencies. In other words, alleles with extremely low and high derived allele frequencies, corresponding to alleles that tend to be very young or very old, respectively, in the sample, will tend to be upweighted because of their low minor allele frequencies. The conceptual analog in the case of branches in a genealogical tree is that the (young) branches near the leaves and the (old) branches near the root will be upweighted in the eGRM (**Figure S1A**). We reasoned that this practice would negatively impact the ability of the eGRM to discern population structure. Structure in humans (and in most species in general) are likely established towards the leaves of the tree, perhaps within the last several hundred generations compared to the coalescent history of the sample, and thus ancient alleles or branches on the tree pre-dating the structure of interests will likely carry little information and instead contribute to the relatedness shared across all individuals (Fan et al. 2022; Zaidi & Mathieson 2020). Indeed, we observed in simulations of a simple two sub-population two-way admixture model (**Figure S2**) that branches connecting two individuals from the same subpopulations tend to be much more recent than branches shared by the two sub-populations (**Figure 2A**). However, in the original weight formulation, 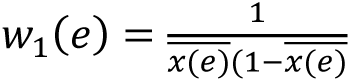, these branches are not up-weighted compared to branches connecting individuals across sub-populations, particularly after accounting for the expected number of mutations on these branches (**Figure 2B**). We thus experimented with different parametric weighting functions (**Figure S3**) and opted to use the weights of the form 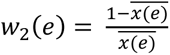 to be effective in up-weighting the more recent past of the genealogical tree (**Figure 2C**). Indeed, the as-eGRM using the updated weight shows clearer contrast between individuals within the same sub-population compared to as-eGRM using the original weights, resulting in clearer demarcation of the two sub-populations on principal components analysis of the as-eGRM (**Figure 2D**).

**Figure 2.**
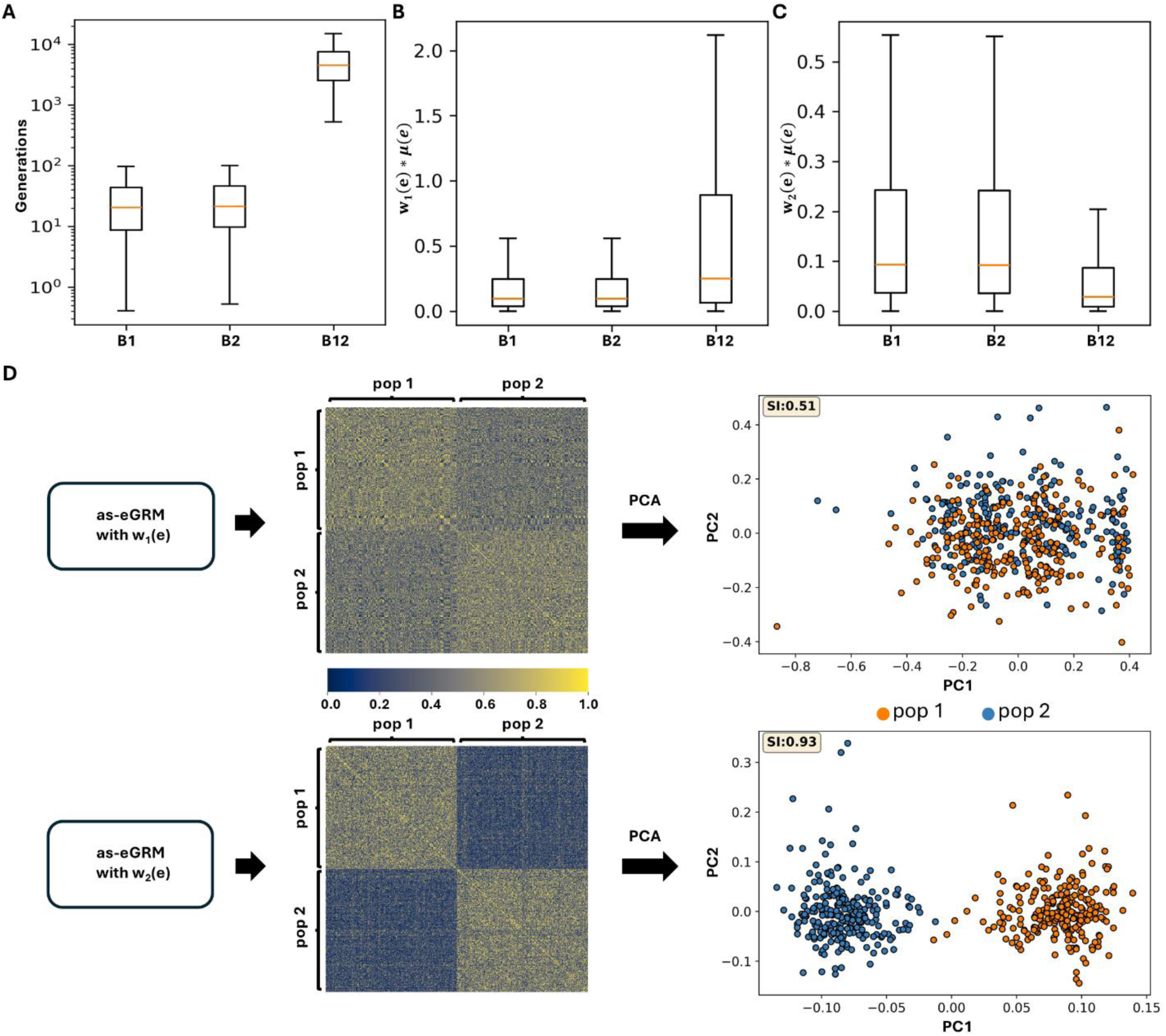
Up-weighting recent branches enhances as-eGRM performance in revealing finer-scale structure in admixed populations. An admixed population with a two-subpopulation (labeled pop1 and pop2) structure was simulated using the model in **Figure S2**. **(A)** Population-common branches are more ancient than the population-specific branches. B1, B2, and B12 represent branches specific to pop1, pop2, and common to both, respectively. Population-specific branches and population-common branches are computationally defined as the branches with more than 80% of the descendants coming from one population (i.e. pop1 or pop2) and the branches with the descendants cover more than 40% of the individuals from each of pop1 and pop2, respectively. **(B)** *w*_1_(*e*) denotes the weighting function used by eGRM. When *μ*(*e*) is weighted by *w*_1_(*e*), population-specific branches are not up-weighted relative to the population-common branches. **(C)** *w*_2_(*e*) denotes the current weighting function used by as-eGRM. When weighted by *w*_2_(*e*), population-specific branches are upweighted when computing the expected relatedness between pairs of individuals because of the greater weight placed on recent branches. **(D)** as-eGRM resulting from the two different weighting functions show different levels of contrast between individuals from each of the sub-populations. as-eGRM using *w*_2_(*e*) results in intra-population relatedness values that are significantly higher than inter-population values, facilitating PCA-based population separation. The as-eGRMs were visualized as heatmaps. To aid in visualization, we rescaled the middle 90% of the as-eGRM values to be within range of 0 to 1 and set the outlier to the boundary values. PCA was applied to the original, untransformed, as-eGRM.

### as-eGRM outperforms alternative methods in extensive simulation

We used a two-split two-way admixed demographic model to simulate an admixed population with structure for evaluating the performance of as-eGRM in revealing fine-scale structure (**Figure 3A**). In this model, there is a first population split 2000 generations ago, separating the orange ancestry (anc2) from the blue ancestry. A second split then occurred at 100 generations ago, creating anc1 population as well as a 3×3 stepping stone model with bi-directional migration with rate 0.01 with neighboring demes to establish a grid-like spatial structure. Finally, 20 generations ago there is a single pulse admixture from anc2 to the 9 demes, with varying proportions (**Figure 3B**). We assessed the performance of as-eGRM using the Separation Index (SI) (Fan et al., 2022), which quantifies the proportion of nearest neighbors belonging to the same subpopulation in the simulated “ground truth” multi-dimensional space. A higher SI indicates better performance. When we applied PCA+UMAP to the canonical GRM from the simulated data, we observed the appearance of approximately 9 demes, though there are clear misclassifications of individuals that are driven by similar ancestry proportions (**Figure 3C**, **Figure S4**; r = -0.43 and -0.54 between ancestry proportions and UMAP1 and UMAP2, respectively). When PCA+UMAP was applied to the eGRM without taking into account local ancestry information, there is again little power to differentiate the structure specific to the blue ancestry (anc1; SI = 0.21). While UMAP applied to the result of AS-MDS or mdPCA showed some improvement (SI = 0.36-0.38) over the result from eGRM, the resolution is limited (**Figure 3C**). In contrast, as-eGRM was able to clearly delineate the 9 demes, completely free from the influence of admixture from anc2 (**Figure 3C**).

**Figure 3.**
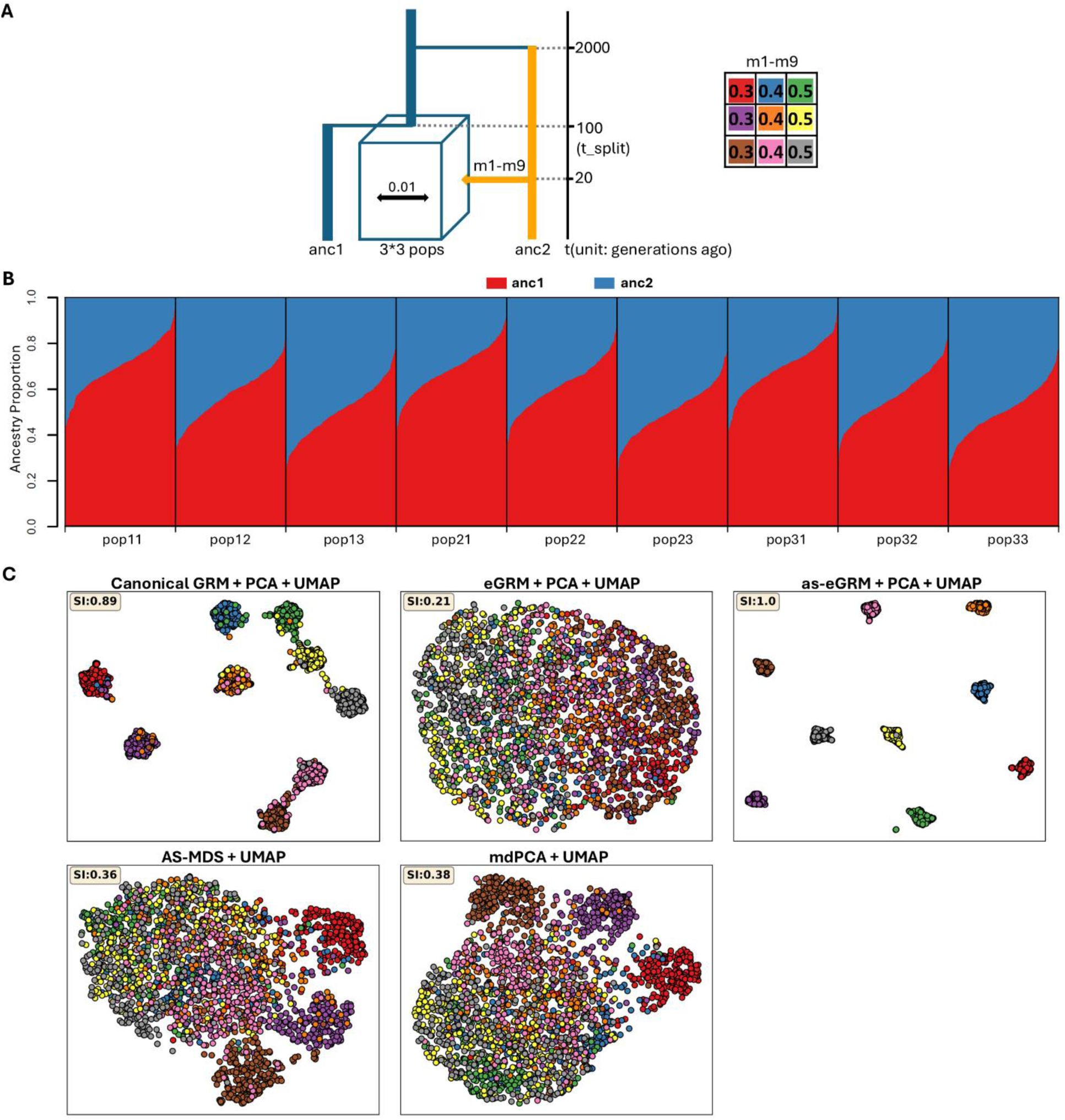
as-eGRM outperforms alternative methods when applied to an admixed population with a grid-like spatial structure. **(A)** The demographic model for simulating an admixed population with a 3*3 grid-like subpopulations structure. *anc1* and *anc2* represent ancestral populations, and were used as the reference for local ancestry inference. *m1*-*m9* specify the proportions of genomic components derived from *anc2* for the individuals in the nine demes, respectively. *t_split* and *t_admix* specify the time the nine subpopulations split and the time the admixture event happened, respectively. Recent migration (rate: 0.01) between neighboring demes has occurred over the last 10 generations **(B)** Ancestry proportions of the individuals in the nine subpopulations, as inferred by RFMix. **(C)** The performance of PCA followed by UMAP applied to the canonical GRM, the eGRM, the as-eGRM, as well as UMAP applied to AS-MDS and mdPCA. 20 PCs were projected down to 2 dimensions by UMAP, as shown in biplots. Data points represent individuals, with colors indicating population membership. Axes for UMAP plots are not labeled as distances are meaningless after UMAP transformation.

We also investigated the impact of different admixture proportions from anc2 as well as the timing of the split to establish the 3×3 grid structure on each method’s ability to discern population structure. Greater admixture from the non-target ancestry would reduce the portion of the genome that are informative for fine-scale structure in the ancestry of interest, and more recent structure would also mean less differentiation among the demes, making fine-scale structure less discernable. We thus conducted additional simulations and evaluations with setting the admixture proportions m1-m9 to 0.2∼0.4 and 0.4∼0.6 and setting structure ages t_split to 50 and 300, separately. As expected, the performance for AS-MDS and mdPCA decreased with increasing admixture proportions from anc2 (**Figure S5**) or more recent onset of the grid-like structure (**Figure S6**) both visually in biplots and by SI. The performance for both ancestry-specific approaches also improved when admixture proportions from anc2 decreased, or when the grid-like structure persisted for longer (**Figure S5, S6;** SI = 0.74-0.86). In all scenarios, as-eGRM consistently outperforms the alternatives, with near perfect delineation of the nine demes (**Figure S5, S6**). The consistently poor performance of eGRM across scenarios highlights the benefits of the modifications implemented in as-eGRM.

We further evaluated as-eGRM on a more realistic three-way admixed Latino demographic history previously fitted from the inferred genealogical trees from array genetic data of Latinos residing in Los Angeles, CA (Fan et al. 2023). We modified this model to include recent population split at 50, 100, or 300 generations ago (**Figure 4A**). Both subpopulations recived same amount of introgression from two other ancestries (10.7% from an “African-like” ancestry and 44.2% from an “European-like” ancestry) at 25 generations ago (**Figure 4B**). Again, as-eGRM outperformed the canonical GRM and eGRM in discerning the population structure in PCA (**Figure 4C**), including scenarios with more recent structure (**Figure S7**).

**Figure 4.**
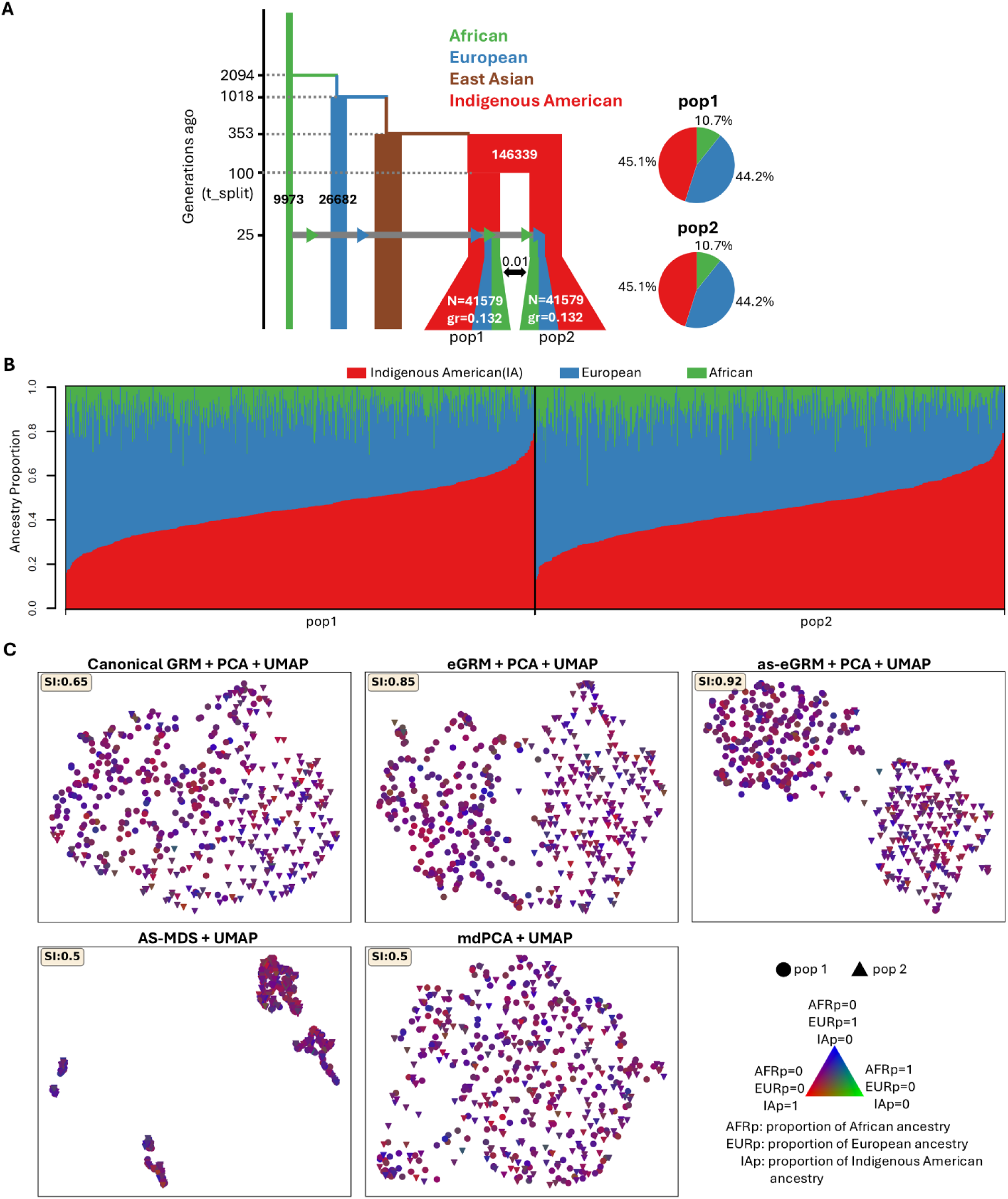
as-eGRM outperforms alternative methods when applied to simulated Latino populations. **(A)** The demographic model for simulating a Latino population with a two-subpopulation structure. Recent migration (rate: 0.01) between the two subpopulations has occurred over the last 10 generations. The model is adapted from (Fan et al. 2023). The ancestral populations African, European, and Indigenous American were used as the reference for local ancestry inference. **(B)** The ancestry proportions of the individuals in the two subpopulations, as inferred by RFMix. **(C)** The performance of PCA followed by UMAP applied to the canonical GRM, eGRM, and as-eGRM, and UMAP applied to AS-MDS and mdPCA, on revealing the two-subpopulation structure. 10 PCs were projected down to two dimensions by UMAP, shown as biplots. Data points represent individuals, with shape and color denoting population membership and ancestry proportions, respectively, as annotated in the lower right corner. In this figure, t_split=100 was used in simulation; see **Figure S7** for the scenarios where t_split = 50 or 300. Axes for UMAP plots are not labeled as distances are meaningless after UMAP transformation.

### as-eGRM outperforms alternative methods in empirical data

We applied as-eGRM to genotyping array data of individuals from Latin America to evaluate its ability to delineate fine-scale population structure in empirical analysis. Many of the Latino individuals have admixed genomes consisting of three predominant continental ancestries: Indigenous American (IA; primarily of South and Central America, Mexico, and the Caribbean islands), European as a result of colonization, and African as a result of slave transport from West Africa (Conomos et al. 2016). We focused on visualizing the fine-scale structure from the IA ancestry component.

We first examined the Latino populations from the Population Architecture using Genomics and Epidemiology (PAGE) study (Wojcik et al. 2019). We take the country of origin as the truth, hypothesizing that different countries across the Central and South America will be correlated with the fine-scale structure within the IA ancestry component. We found that PCA or PCA followed by UMAP approaches generally can discern the population structure in this dataset, though PCA based on the canonical GRM or the eGRM appears to be driven by ancestry proportions (**Figure S8,** left column; r = -0.79 and -0.73 for PC1 of canonical GRM and eGRM, respectively; r = -0.12 and 0.21 for PC2 of canonical GRM and eGRM, respectively). All ancestry-specific methods (i.e. AS-MDS, mdPCA, and as-eGRM) outperform PCA on canonical GRM and eGRM and are relatively free from bias by global ancestry (r = -0.05 to 0.33 across methods). Based on the separation index, as-eGRM produced slightly more accurate clustering (based on grouping individuals from different country of origin), though the differences are small (SI = 0.85 for as-eGRM vs. 0.81 or 0.82 for AS-MDS and mdPCA; **Figure S8**). All methods performed similarly by separation index when UMAP is applied to the PCs (**Figure S8**). The general ability for each method to delineate the population structure may be due to the relatively high level of Indigenous American ancestry in this sample (**Figure S9**). The fact that PCA based on the canonical GRM or eGRM can also somewhat elucidate the IA-specific structure suggests that the IA component across individuals in this dataset may be sufficiently differentiated and correlated with country of origin.

We then studied the Latino population from the Hispanic Community Health Study/Study of Latinos (HCHS/SOL). Previous studies applied the AS-MDS to the HCHS/SOL data and identified fine-scale structure within the IA ancestry that is consistent with grandparental country of origin (Browning et al. 2016). Given that individuals with low levels of IA ancestry will have limited genetic data after masking, thereby adding noise to the PCA, previous studies also restricted their analysis to only individuals with at least 50% of their genomes derived from the ancestry of interest. Indeed, when we restricted our analysis to the subset of individuals with estimated IA ancestry > 0.5 (across all recruitment center), all ancestry-specific methods were able to delineate the population structure better than PCA on the canonical GRM and eGRM (SI = 0.88-0.96 vs. 0.65 and 0.67 on canonical GRM and eGRM, respectively; **Figure S10,** left column). When examining the distribution of IA ancestry across individuals, all methods except the as-eGRM show substantial correlation with IA ancestry on either of the first two PCs (**Figure S10,** left column; |Pearson’s correlation r| = 0.72-0.78). Applying UMAP on the top 50 PCs to collapse them down to 2-dimensions further improved the delineation of population structure for all methods (SI = 0.84 to 0.97 across all methods; **Figure S10**, right column). Consistent with previous report (Browning et al. 2016), we observed clearly 3 to 4 clusters in this dataset, corresponding to Latinos from northern part of Central America (Mexico), southern part of Central America (Costa Rica, El Salvador, Guatemala, Honduras, and Nicaragua), and Southern America (Argentina, Colombia, Ecuador, and Peru).

However, when we applied each method to HCHS/SOL data spanning the entire spectrum of IA ancestry (**Figure S11**), the advantage from as-eGRM become apparent. In this most inclusive scenario, we found that neither of the frequency-based approach (AS-MDS and mdPCA) nor the non-ancestry-specific approach (PCA on canonical GRM and eGRM) could appropriately delineate the structure as defined by grandparental country of origin (**Figure 5**) that was more apparent when only analyzing the subset of individuals with high IA ancestry (**Figure S10**). Any pattern that was discernable from PCA were strongly correlated with global ancestry, particularly the European ancestry (**Figure 5**, left column; |Pearson’s correlation r| = 0.7-0.86). In contrast, as-eGRM significantly outperformed all alternatives; it showed clearer separation by major grandparental country of origin in PCA, which is not confounded by proportion of global ancestry (**Figure 5**). Applying UMAP on the top 50 PCs somewhat improved the clustering (**Figure 5**, right column). However, while the expected clusters based on northern Central America, southern Central America, and South America may start to separate in analysis using the canonical GRM, eGRM, or AS-MDS and mdPCA, they are far from the clean distinct clusters when as-eGRM was used.

**Figure 5.**
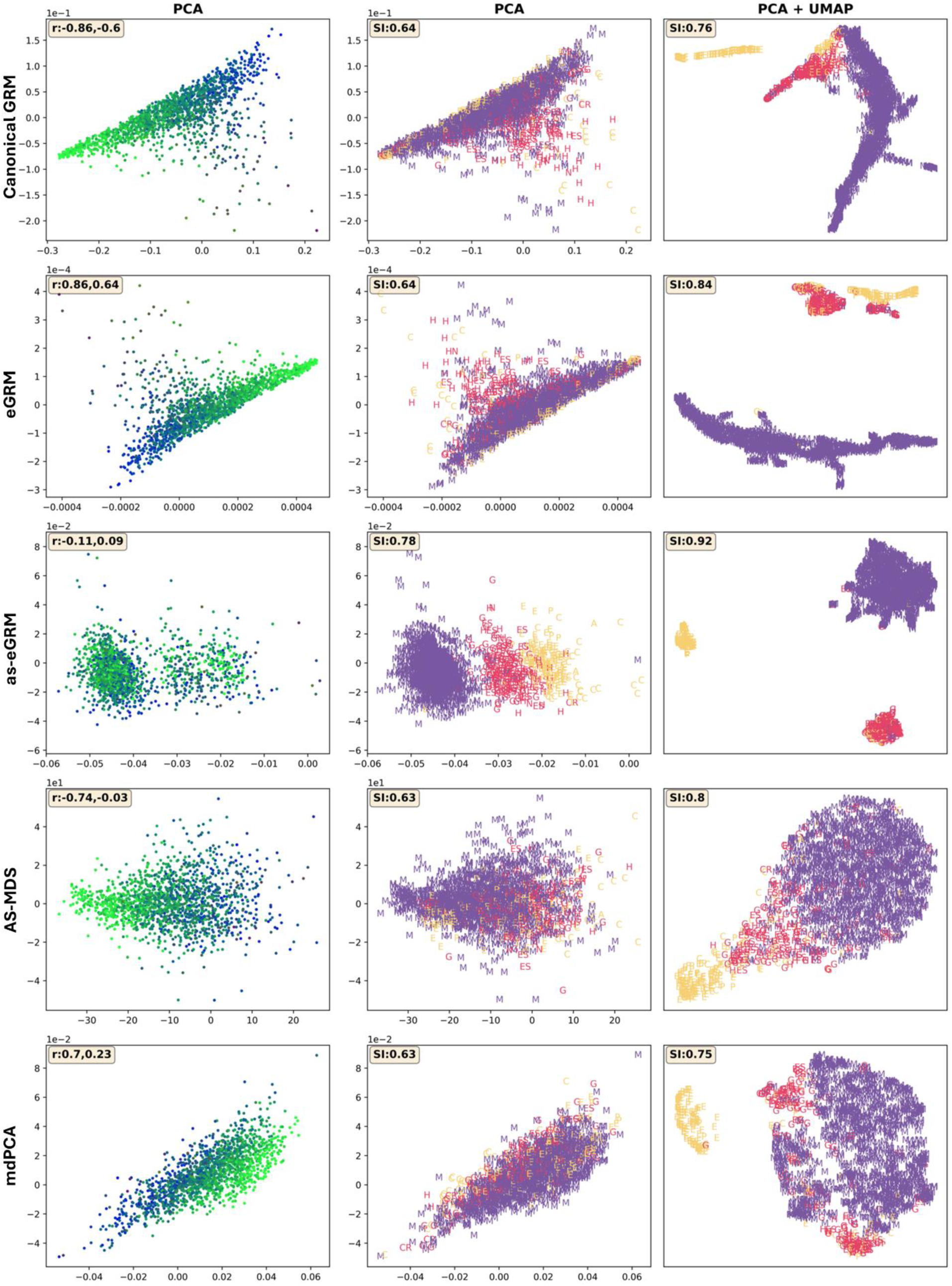
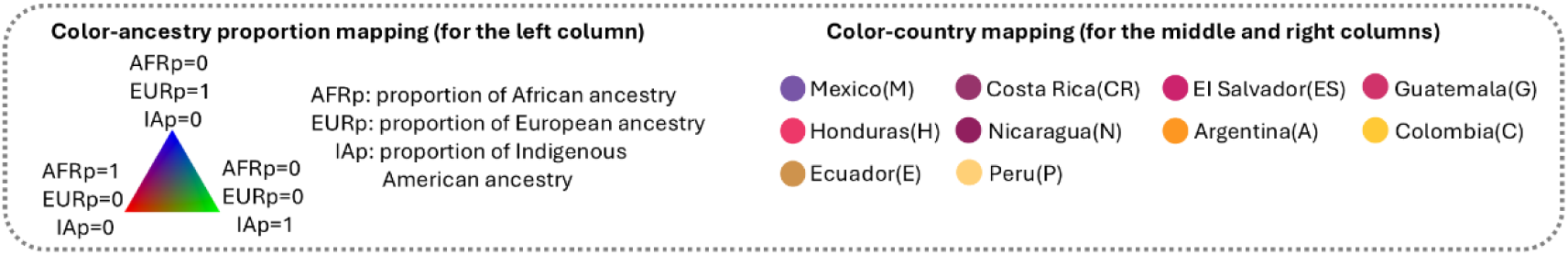
as-eGRM outperforms alternative methods in revealing the Indigenous American ancestry-specific structure in the Hispanic/Latino population using HCHS/SOL data. Analysis focused on 1671 individuals from the Chicago recruitment site only but spanned the entire spectrum of ancestry proportions. Plots showed PCA or PCA+UMAP results (column annotations) of each analytical approach (row annotations). Points represent individuals, colored by ancestry proportions (left column) or grandparental country of origin (middle and right columns, see bottom box for annotation). The r in the left upper corner of the left column represents the Pearson correlation coefficients between the proportions of Indigenous American ancestry and PC1 (the first number) or PC2 (the second number), respectively. The Separation Index (SI) in the left upper corner of the right two columns is calculated using grandparental country of origin as (presumed) true labels. Axes for UMAP plots are not labeled as distances are meaningless after UMAP transformation.

## Discussion

In this study, we introduced as-eGRM, a framework that leverages genealogical trees and local ancestry information to reveal ancestry-specific structures in admixed populations. The key advancements of as-eGRM include defining ancestry-specific pairwise relatedness between individuals based on genealogical trees and local ancestry callsets across the genome accounting for missing data (due to masked non-target ancestry), as well as a modified weighting of branch on the trees to up-weight recent branches more informative of recent population structure. Through extensive evaluation using multiple simulated and empirical datasets, we demonstrated that as-eGRM consistently outperforms alternative metrics or methods in revealing ancestry-specific population structure across various demographic scenarios and missing data proportions.

In this study, we opted to illustrate the power of our method using individuals from Latin America from the HCHS/SOL dataset as the empirical example. Latinos are known to exhibit substantial heterogeneity in the distribution of their genetic ancestry across Latin America (Price et al. 2007; Gravel et al. 2013), or even across geographical locations within the United States (Bryc et al. 2015). Yet, Latinos across geographical space tend to be aggregated for analysis. Combined with heterogeneous exposure through the environment, such aggregation has been shown to mask differences in the phenotype distribution, efficacy of polygenic risk score, and evaluation of interaction between ancestry and environment (Sharma et al. 2024). These differences can become more apparent in a stratified analysis even if just based on the self-reported ethnicity or country of origin (Sharma et al. 2024). Furthermore, the definition of the Indigenous American component of genetic ancestry tend to also aggregate putative reference individuals across Latin America, again masking the allele frequencies difference that arise in ancestral Indigenous American populations across the Americas due to their unique migration history (Skoglund & Reich 2016; Scheib et al. 2018). Indeed, previous assessment of population structure in Latin America have also shown the fine-scale differentiation within a broad umbrella term of Indigenous American ancestry (Moreno-Estrada et al. 2013; Sohail et al. 2023). The as-eGRM thus stands to improve these investigations leveraging the genetic linkage information. By providing a more nuanced view of genetic variation within (somewhat arbitrarily defined) ancestry components, as-eGRM help re-define or re-interpret genetic ancestry, improve our understanding of population history, and may lead to more robust and interpretable findings in studies of diseases and complex traits. This advancement could pave the way for more precise and equitable genetic research, ultimately contributing to better health outcomes for diverse populations.

We also opted to utilize UMAP to complement in exploring the population structure of our simulated and empirical datasets. There has been recently well-known discussion on social media regarding PCA and UMAP, in the context of their applications to represent the genetic diversity of the All-of-Us cohort (The All of Us Research Program Genomics Investigators et al. 2024). While the majority of the criticism (Pachter 2024) centered on the conflation of genetic ancestry and self-reported race and ethnicity through questionable use of color and labels, UMAP was also suggested to contribute towards forcing a discrete nature of genetic diversity in an inherently continuous space (as visualized by PCA plots). Indeed, while the admixture process in humans is modeled as an inherently linear process, UMAP does not preserve the distances in its transformation but instead accentuate the distinctiveness of the majority subgroups. However, both PCA and UMAP, when applied to genetic data, are dimensional reduction approaches to reduce the high-dimensional genetic data down to visualizable 2- or 3-dimensions for exploratory analysis. The representation of the genetic data will not be loss-less through any form of dimensional reduction techniques, and the appropriate usage may depend on the context. The appropriate use of UMAP on human data is continually being explore (Diaz-Papkovich et al. 2023), and it may be more suitable in the context of exploring isolated islands where significant drift may occur, for instance (Ioannidis et al. 2021). In our context, simulated data assumed a discrete nature (e.g. the 9-deme model; **Figure 3**) and UMAP could be more powerful in identifying these clusters. Similarly, in our empirical application, we targeted the Indigenous American ancestry and assumed that the fine-scale structures of interests are more discrete in nature. Such assumption is made whenever one operates under a generally discrete view of genetic ancestry (when reference ancestral populations are presumed when modeling admixture history, for instance). This may or may not reflect the reality, but we note that UMAP is applied as one potential approach to explore the data and generate additional hypothesis of the history of these populations, to complement the visualization through PCA, which we also show.

On a technical level, we found that population structure analysis based on the as-eGRM excels over previous methods (AS-MDS or mdPCA) when the proportion of admixture from non-target ancestry is high. For instance, as the non-target ancestry increased from 0.2-0.4 to 0.4-0.6 in the nine-deme stepping stone model (**Figure S5**), mdPCA progressively performed worse in elucidating population structure (SI = 0.74 to 0.15), while as-eGRM maintained sensitivity (**Figure S5**). This may also underlie the observation that AS-MDS and mdPCA performed comparably to as-eGRM on the PAGE-Latin American dataset (**Figure S8**; mean IA ancestry proportion = 0.68) or the HCHS/SOL data when filtering on individuals with estimated IA ancestry > 0.5 (**Figure S10**). One reason for this observation is the impact due to missing data. As admixture proportions from non-target ancestry increases, the proportion of the genomes between a pair of individuals that are not masked by AS-MDS or mdPCA decreases, reducing the information available to compute genetic similarity between the pair. Similar issue with pervasive missingness in the GRM had been discussed in literature, particularly when using data from ultra-low coverage sequencing data or ancient DNA (aDNA). Common approaches to deal with missingness when inferring population structure includes filtering of individuals with high missingness or impute the missingness by mean genotype values (Arteaga & Ferrer 2002; Patterson et al. 2006; Galinsky et al. 2016; Abraham et al. 2017), though both approaches could introduce bias in population structure inference. Other approaches, such as those based on an expectation-maximization algorithm to iteratively impute frequency of missing genotypes (Meisner et al. 2021), or based on matrix denoising techniques and truncated SVD as used by mdPCA, have also been proposed to deal with the non-random missingness in the data. In our as-eGRM framework, we did not explicitly deal with missingness in the construction of as-eGRM; we also simply ignored the regions of genome between pairs of individuals where one or both individuals have non-target ancestries. Indeed, we found that the variance in our estimates of ancestry-specific relatedness to be relatively small, oftentimes one order of magnitude lower than the estimates themselves even when missingness is around 90% (**Figure S12**). While our approach appears to be robust to increased admixture proportions, its ability to elucidate population structure may still suffer when investigating structure within a minor ancestry component, or when ancestry segments are not randomly distributed in the genome (*e.g.* in presence of adaptive introgression). We would also expect the variance of the relatedness estimates to be larger if the ARG reconstruction is less accurate, or if less genetic information is available for ARG reconstruction (*e.g.* array genotypes were used). Therefore, future improvements may focus on evaluating and implementing approaches to ensure robustness across the spectrum of missing information.

The current implementation of as-eGRM has some limitations and future direction for improvement. First, we found that up-weighting recent branches is crucial for revealing contemporary fine-scale structures. This finding suggests that selectively weighting of branches from different parts of the trees could enable the detection of structures from specific time periods. While our current approach empirically determines the weighting function for recent branches, future research should explore systematic methods to derive optimal weighting functions for both recent and temporally specific structures. Second, as-eGRM’s reliance on ARG-reconstruction makes it computationally intensive for datasets exceeding a few thousand individuals. ARG-reconstruction methods scalable to biobank level data are available (Wohns et al. 2022; Zhang et al. 2023), though its accuracy can still be improved (Y. C. Brandt et al. 2022; Fan et al. 2022; Peng et al. 2024). We chose to use Relate as the best combination of accuracy and scalability and also expect that rapid advances in ARG-reconstruction methods will likely improve both the accuracy and scalability, which will benefit the as-eGRM framework. Third, our method cannot yet be applied to aDNA data, as the quality of aDNA data cannot yet be used in local ancestry inference (as the target or the reference), and its incorporation into the ARG is still in development. Nevertheless, their incorporation into population structure analysis may be illuminating for both understanding the history of a modern admixed population or in interpreting or re-defining the ancestries of an admixed individual. For the time being, allelic-based approaches for incorporating aDNA may still be most reliable. In fact, the explicit reliance of defining a high-quality reference panel and inference of discrete local ancestry labels is a strong limitation of the current approach. While it may be clearer to define ancestral populations for continentally admixed populations, the notion of ancestry is complicated by both geographical location and temporal reference (Mathieson & Scally 2020). Genealogical trees potentially enable a continuous view of population structure and ancestry across time, moving beyond traditional discrete ancestry classifications. Therefore, future development may also move towards a more fluid definition of ancestries and investigate the population structure at multiple levels within a cross section of time.

## Data and code availability

We have implemented the algorithms related to as-eGRM in a python package, asegrm, which is publicly available in PyPI. Documentation of this package as well as the codes for reproducing the analyses in this study can be found on its GitHub page (https://github.com/jitang-github/asegrm).

## Author Contributions

C.W.K.C. conceived of and designed the study. J.T. implemented the method and performed the analysis. J.T. and C.W.K.C. interpreted the data. J.T. and C.W.K.C. wrote the paper.

## Acknowledgement

We would like to thank Bryan Dinh and Jalen Langie for providing curated datasets and tools that made the analysis of this study feasible. We would also like to thank John Novembre, Arun Durvasula, Shaila Musharoff and other attendees of the 2024 American Society of Human Genetics annual conference for encouragement and discussion of this method. Research reported in this publication was supported by the National Institute of General Medical Sciences (NIGMS) of the National Institute of Health, under award number R35GM142783 (to C.W.K.C.). Computation for this work is supported by USC’s Center for Advanced Research Computing (https://carc.usc.edu/).

## Declaration of interests

The authors declare no competing interests.

## Supplementary Materials

**Figure S1.**
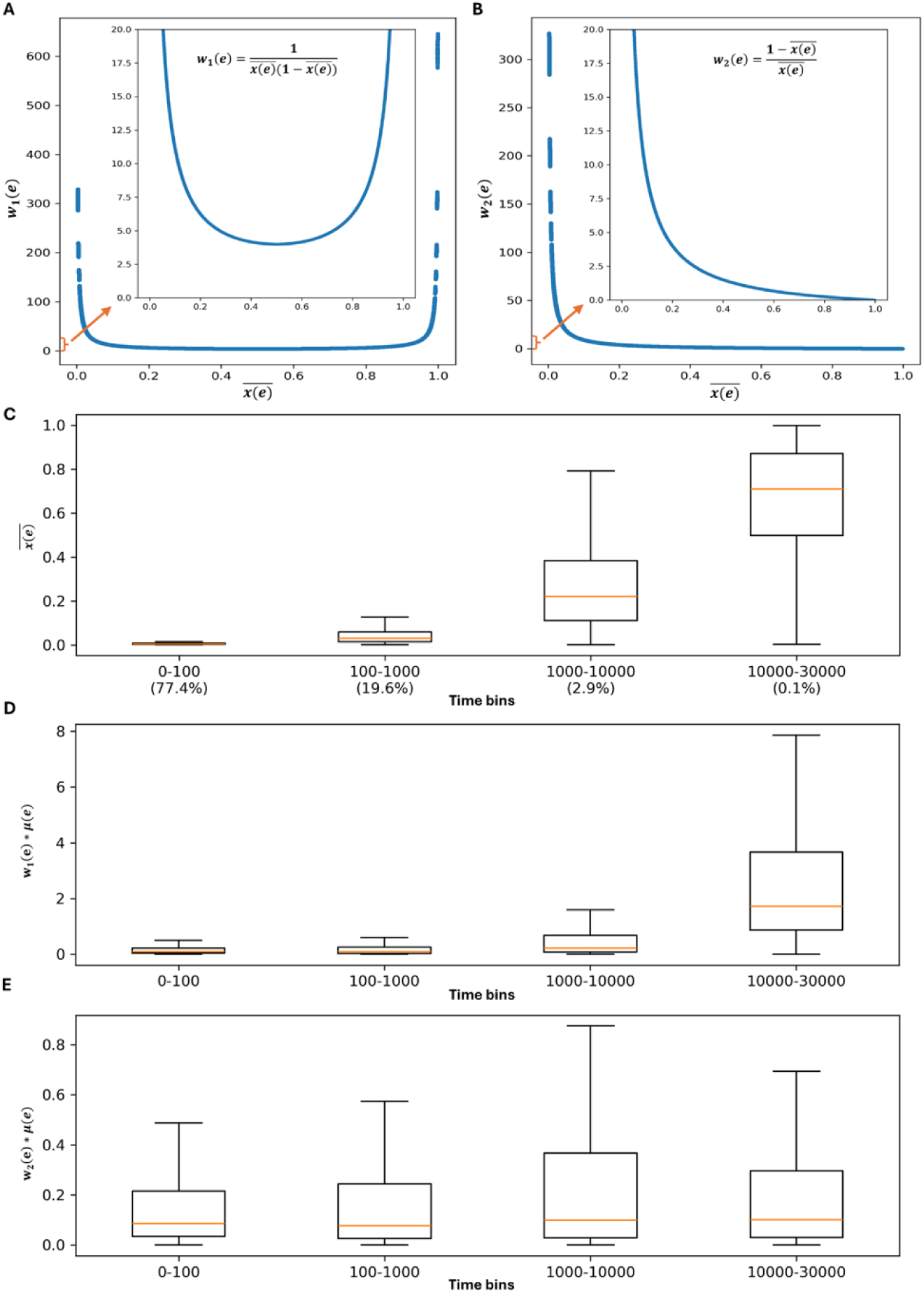
The comparison of the weighting functions used by eGRM and as-eGRM. (A) The weighting function *w*_1_(*e*) used by eGRM up-weights the branches with a low or high 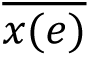, which represents the proportion of the descendants under branch *e* in all descendants. Inset shows the same function but with y-axis capped at 20. (B) The weighting function *w*_2_(*e*) used by as-eGRM up-weights the branches with a low 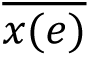. Inset shows the same function but with y-axis capped at 20. (C) In a simulation of 500 individuals over a 100Mb region based on the demographic history of **Figure S2**, we stratified all branches *e* into four time bins. The more ancient time bins tend to have higher value of 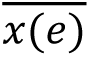. (A-C) indicate that *w*_1_(*e*) up-weighs both recent and ancient branches, while *w*_2_(*e*) up-weights only recent branches. (D) Multiplied by *μ*(*e*), the expected number of mutations occurring on branch *e*, to account for the expected number of mutations given the branch length, *w*_1_(*e*) assigns relatively bigger values to more ancient branches. (E) But *w*_2_(*e*), on the other hand, would assigns comparable values to different ages of branches in the same construct.

**Figure S2.**
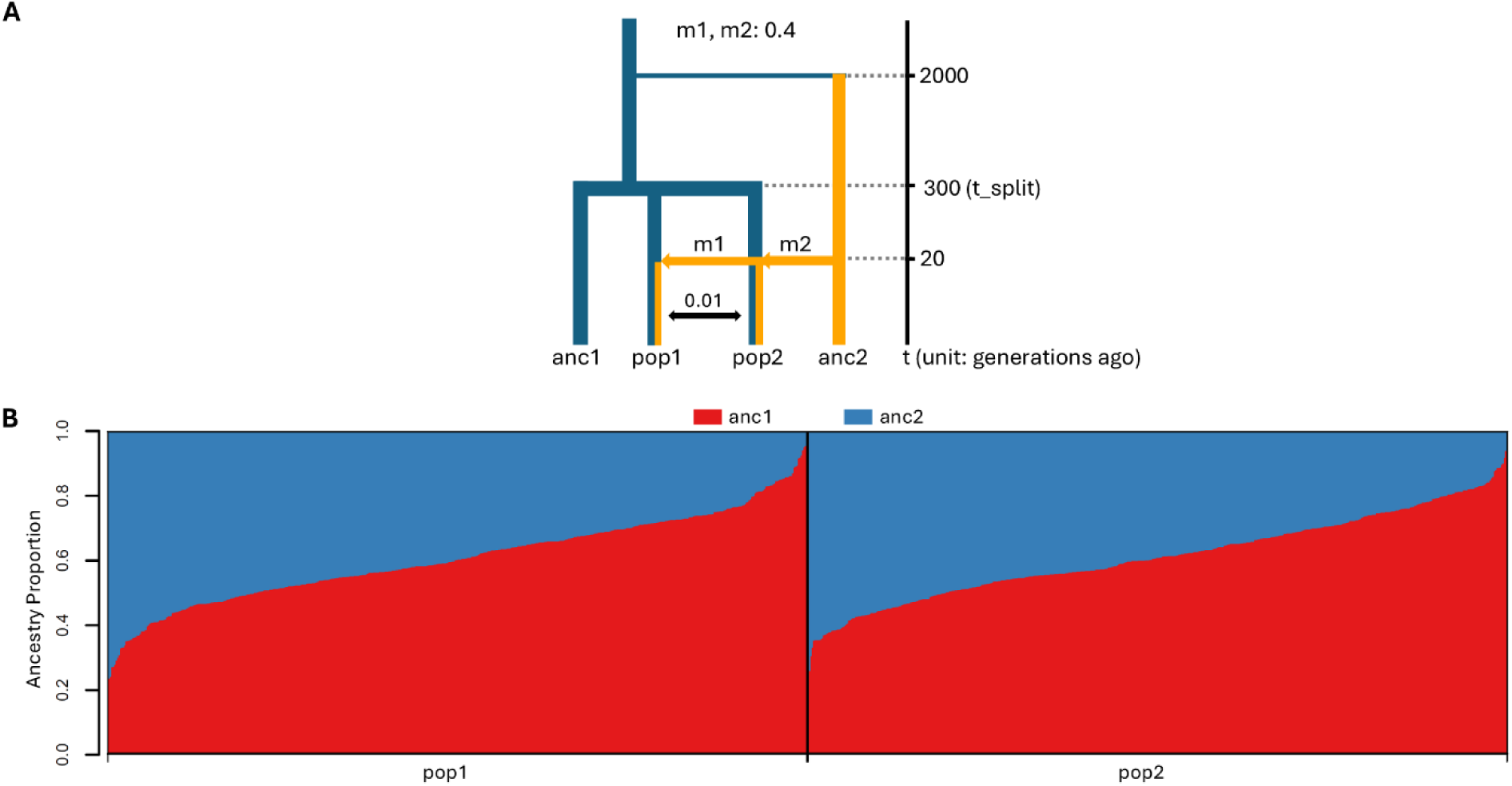
A two-population two-way admixed demography. This simulated scenario was used to explore the effect of different weighting functions in computing the ancestry-specific pairwise relatedness. (A) The demographic model for simulating an admixed population with a two-subpopulation structure. *anc1* and *anc2* represent ancestral populations, and were used as the reference for local ancestry inference. *m1* and *m2* specify the proportions of genomic components from *anc2* for the individuals in *pop1* and *pop2*, respectively. *t_split* and *t_admix* denote the time of *pop1* and *pop2* splitting and the admixture event, respectively. Recent migration (rate: 0.01) between pop1 and pop2 has occurred over the last 10 generations. (B) The ancestry proportions of the individuals in the two sub populations, as inferred by RFMix.

**Figure S3.**
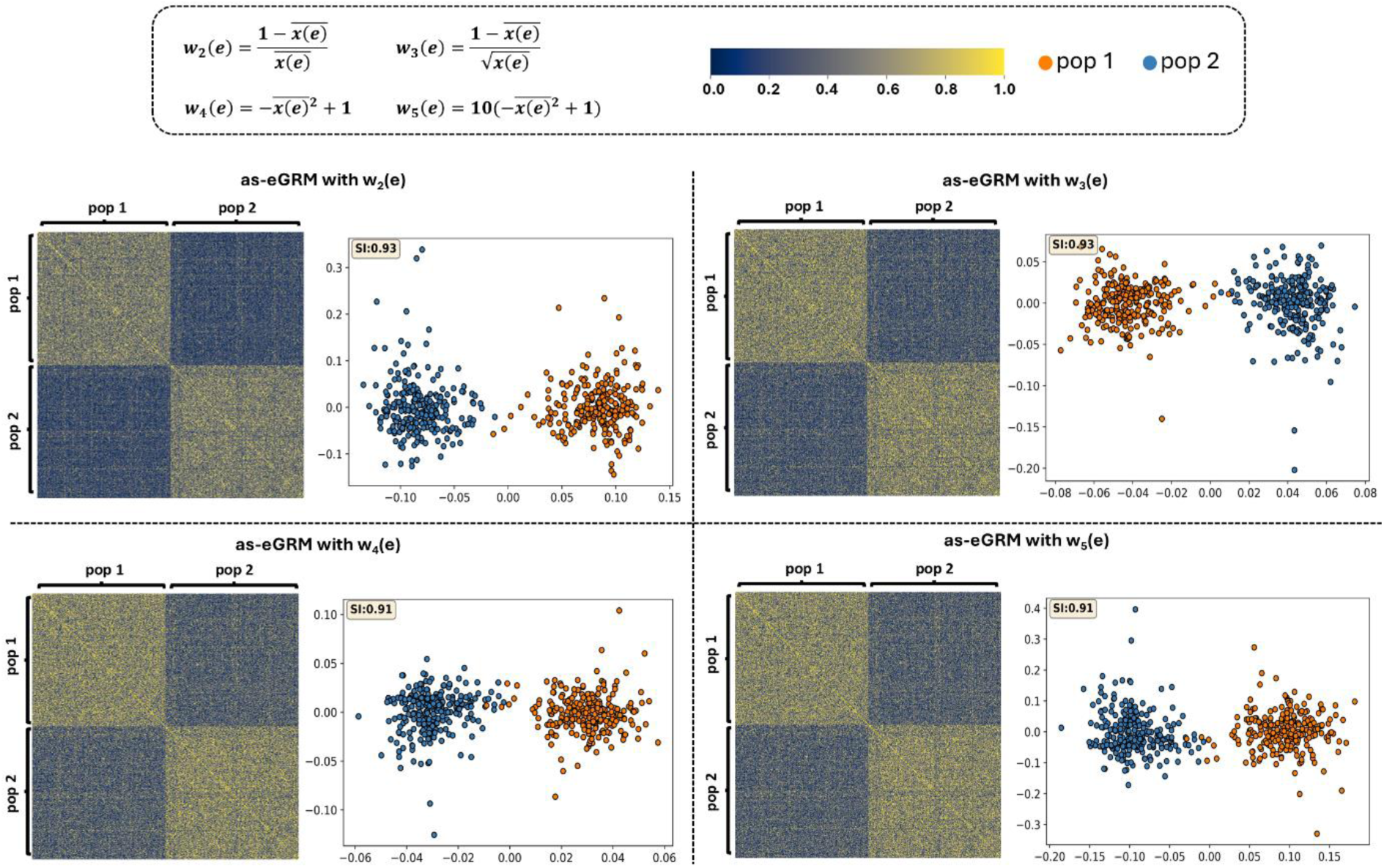
The performance of the candidate weighting functions for up-weighting recent branches. In order to up-weight the recent branches of each genealogical tree to accentuate the recent structure, we searched for a function that increases monotonically as the input branch age decreased. We empirically tried multiple functions with different monotonically increasing slopes, computed and visualized the as-eGRM based on the simulated demography from **Figure S2**, and applied the PCA on the as-eGRM to assess the performance in separating the two sub-populations. The as-eGRMs were visualized as heatmaps. To aid in visualization, we rescaled the middle 90% of the as-eGRM values to be within range of 0 to 1 and set the outlier to the boundary values. PCA was applied to the original, untransformed, as-eGRM. The scatter plots show the top 2 PCs. Based on the performance of these functions, we chose the function *w*_2_(*e*) in this study.

**Figure S4.**
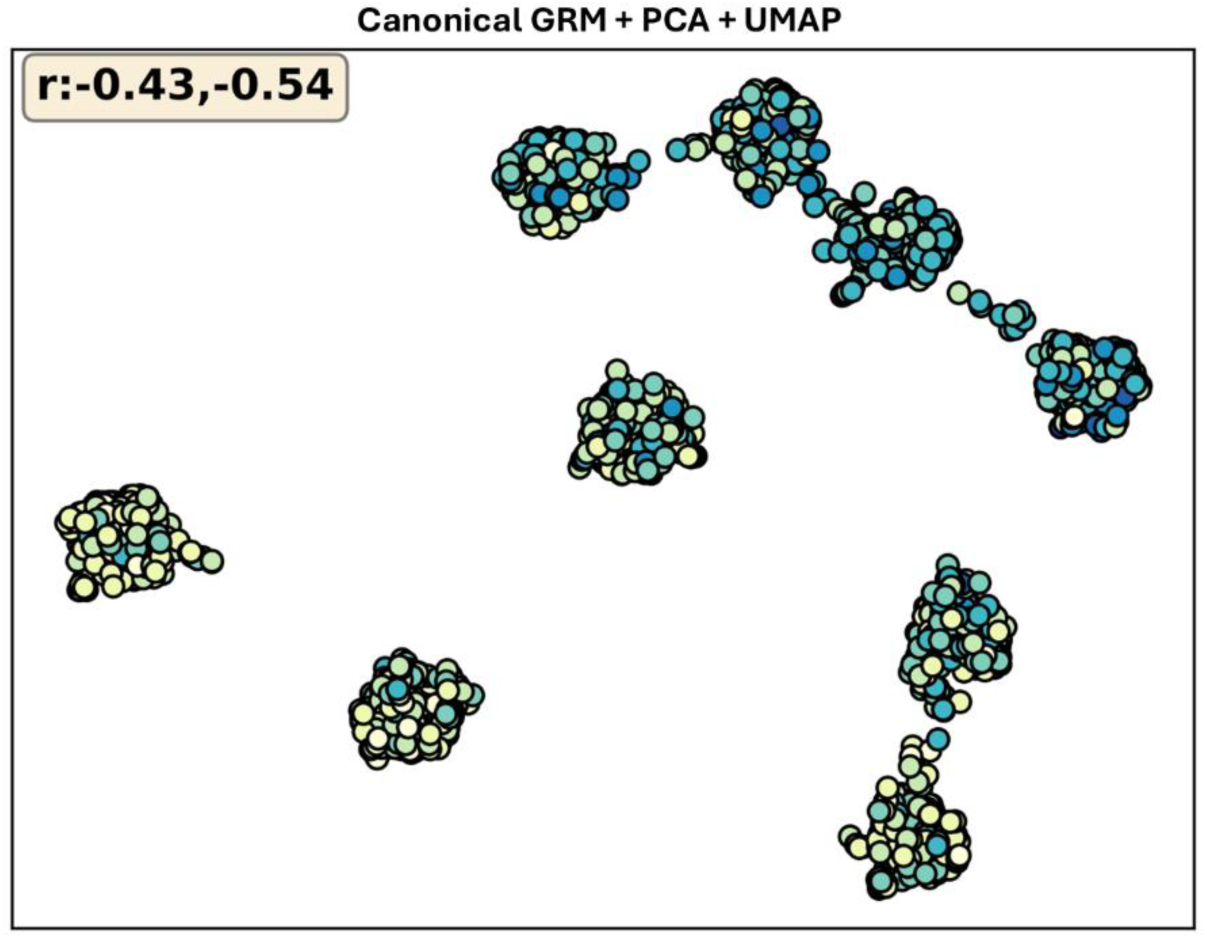
The distribution of the individuals in the PCA+UMAP applied to the canonical GRM is driven by ancestry proportions. 20 PCs were projected down to 2 dimensions by UMAP, as shown in the biplot. Data points represent individuals, with colors indicating the ancestry proportion of the population targeted for investigation. The r in the left upper corner represents the Spearman’s rank order correlation coefficients between the ancestry proportions of the population targeted for investigation and UMAP1 (the first number) or UMAP2 (the second number), respectively. These populations were simulated using the demographic model in Figure 3A. Axes for UMAP plots are not labeled as distances are meaningless after UMAP transformation.

**Figure S5.**
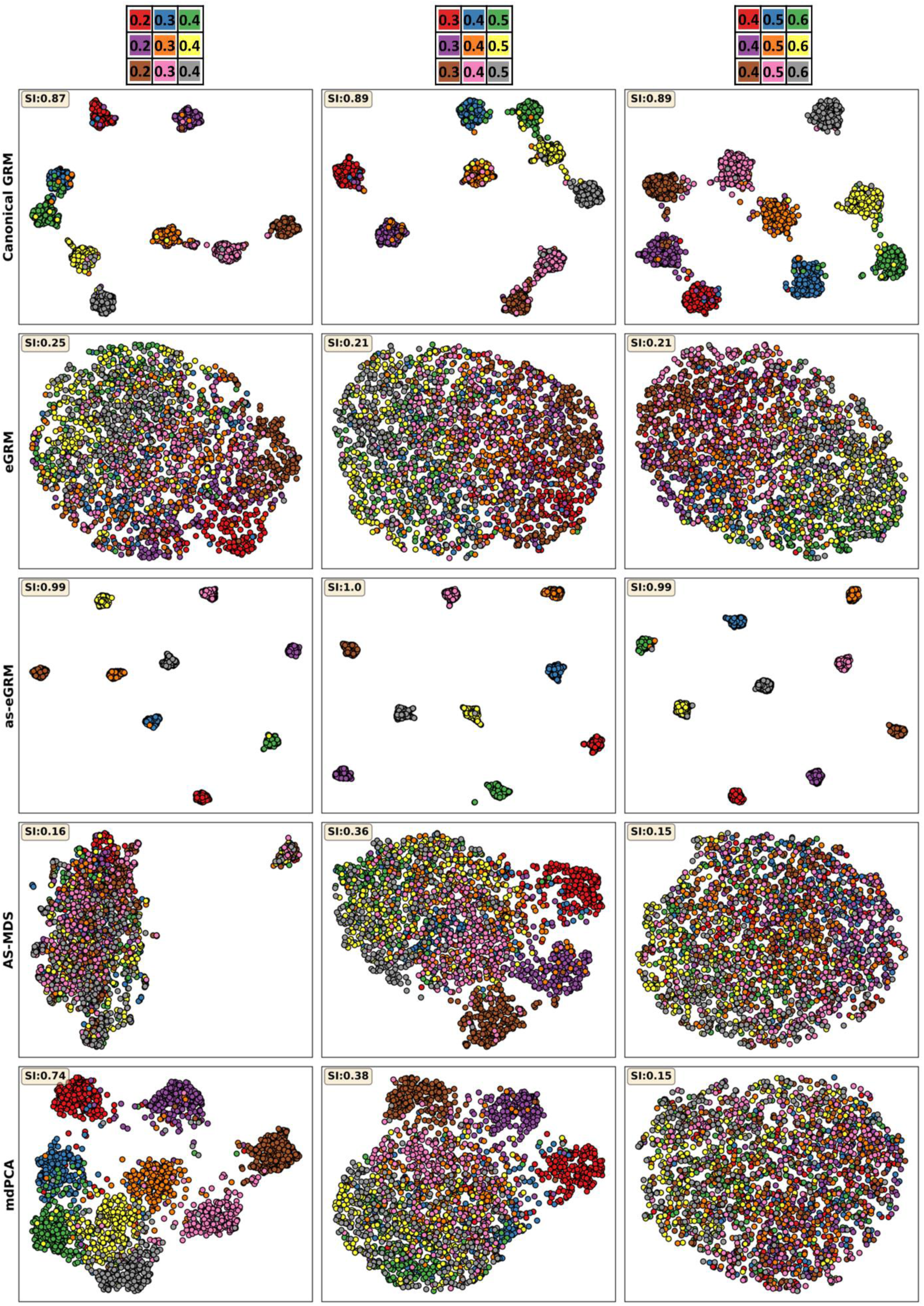
as-eGRM outperforms the alternatives when applied to an admixed population with a grid-like spatial structure across different admixture proportions. 20 PCs were projected down to two dimensions by UMAP, shown as biplots. Data points represent individuals, with colors indicating population membership. These populations were simulated using the demographic model in Figure 3A with the admixture proportions set to the values annotated by the grids on the top row. The other demographic parameters were kept the same as in Figure 3A. Axes for UMAP plots are not labeled as distances are meaningless after UMAP transformation.

**Figure S6.**
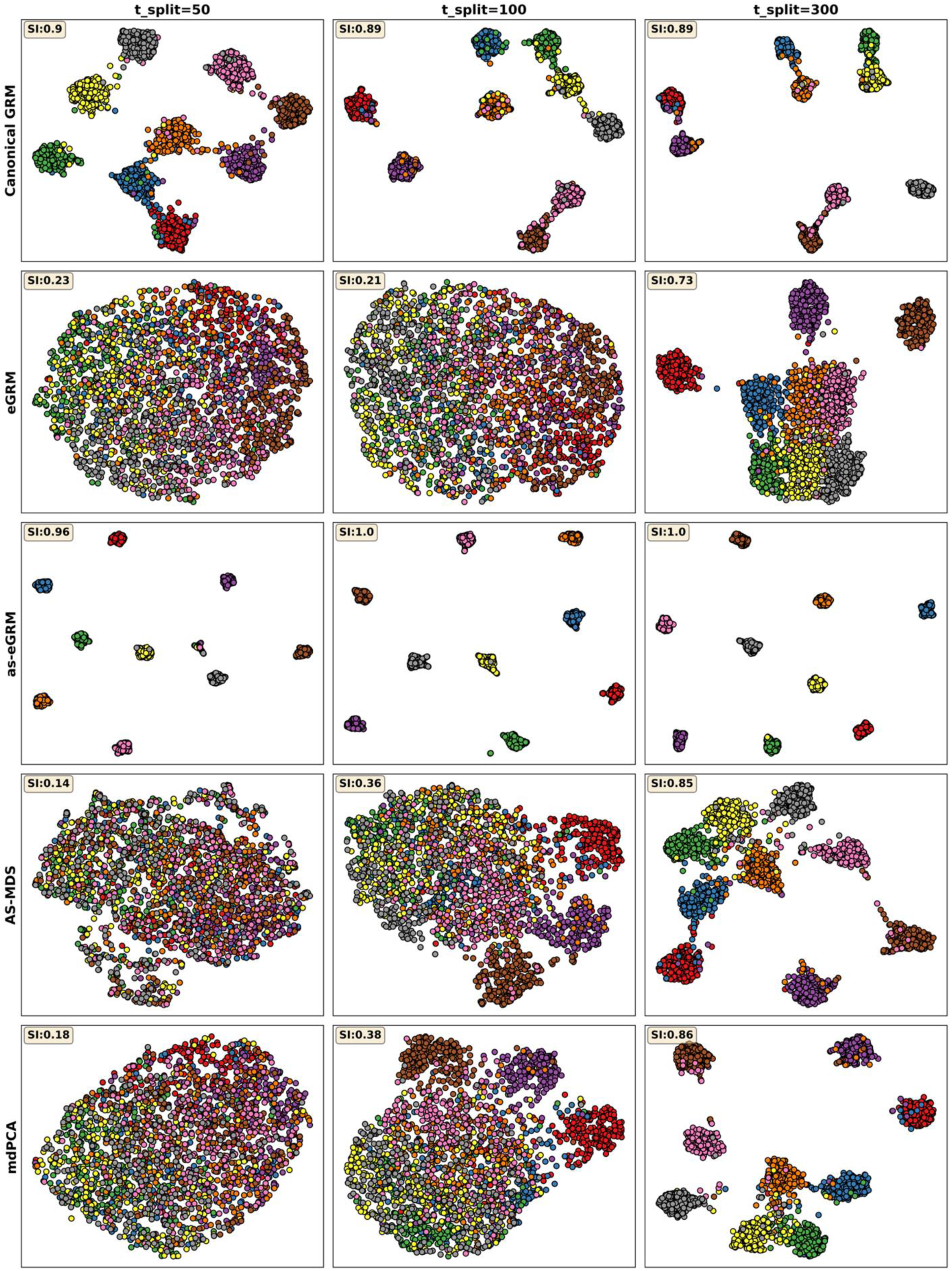
as-eGRM outperforms the alternatives when applied to an admixed population with a grid-like spatial structure across different structure ages. 20 PCs were projected down to two dimensions by UMAP, shown as biplots. Data points represent individuals, with colors indicating population membership. The populations were simulated by the model in (Fig. 3A) with the structure ages set to the values annotated by the column names. The other demographic parameters are specified in (Fig. 3A). Axes for UMAP plots are not labeled as distances are meaningless after UMAP transformation.

**Figure S7.**
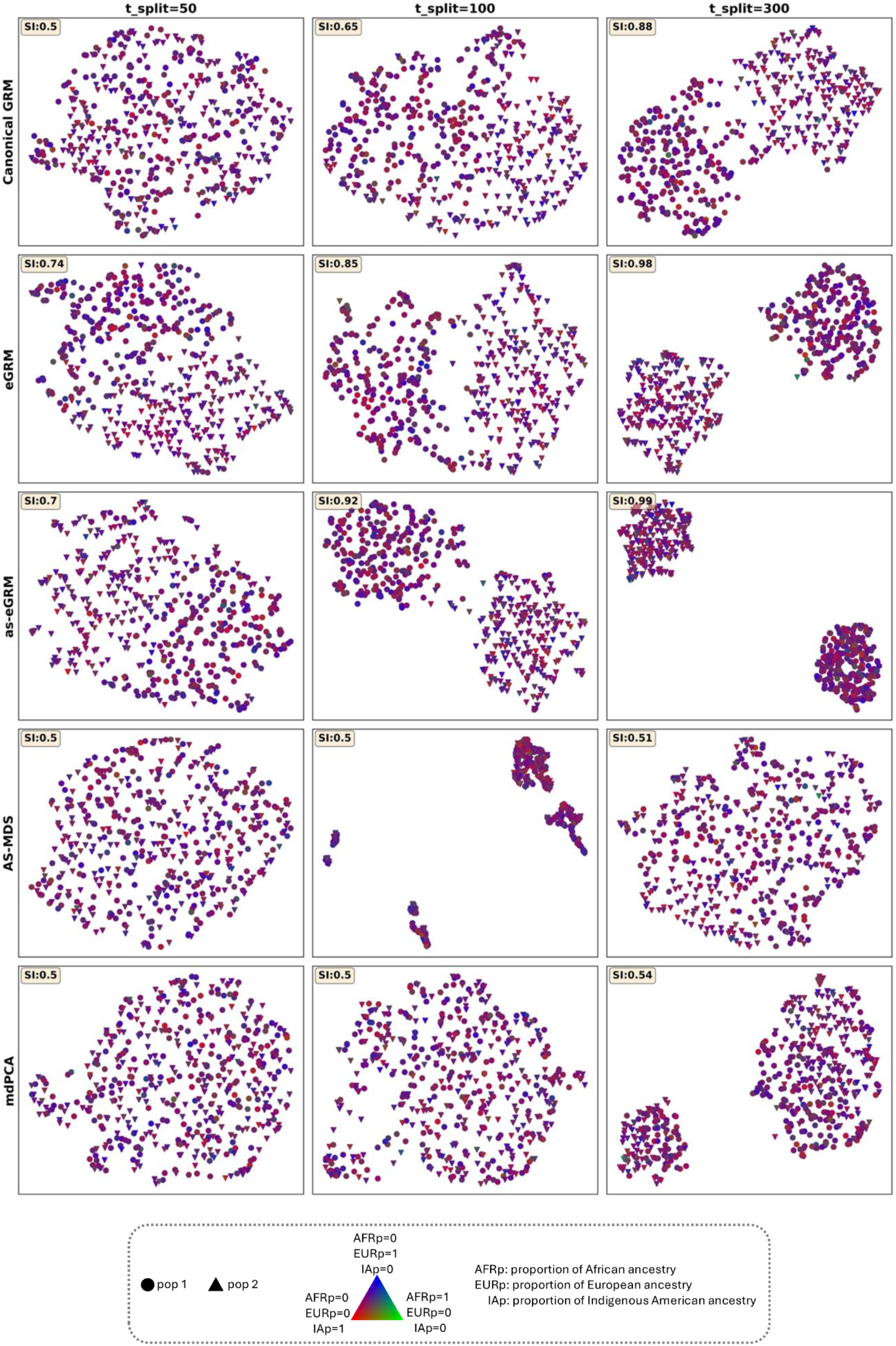
as-eGRM outperforms the alternatives when applied to a simulated Latino population with a two-subpopulation structure across different structure ages. 10 PCs were projected down to two dimensions by UMAP, shown as biplots. Data points represent individuals, with colors indicating ancestry proportions based on the key, and shape of the symbol indicating population membership, as annotated in the bottom box. The populations were simulated by the model in Figure 4A with the timing of the onset of structure set to the values annotated by the column names. The other demographic parameters were kept fixed to that in Figure 4A. Axes for UMAP plots are not labeled as distances are meaningless after UMAP transformation.

**Figure S8.**
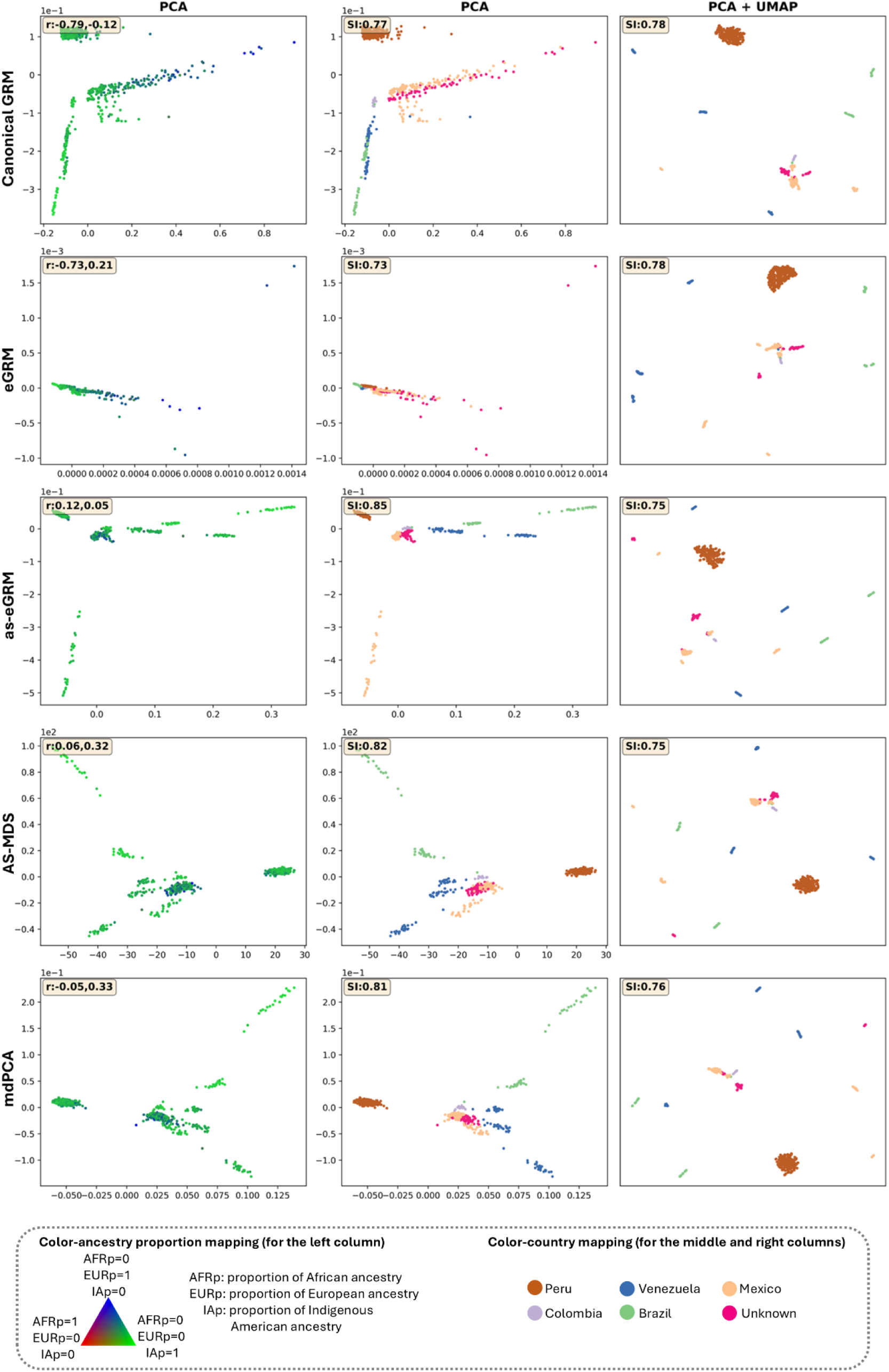
as-eGRM outperforms alternative methods on revealing the Indigenous American ancestry-specific structure in Latin America population using the PAGE data. PCA or PCA+UMAP result (column annotations) for each method (row annotations) when applied to the PAGE global reference panel. Points represent individuals, colored by ancestry proportions (left column) or country of origin (middle and right columns, see bottom box for annotation). The r in the left upper corner of the left column represents the Pearson correlation coefficients between the proportions of Indigenous American ancestry and PC1 (the first number) or PC2 (the second number), respectively. The Separation index (SI) in the left upper corner is calculated assuming the self-reported country of origin as the true labels. Axes for UMAP plots are not labeled as distances are meaningless after UMAP transformation.

**Figure S9.**
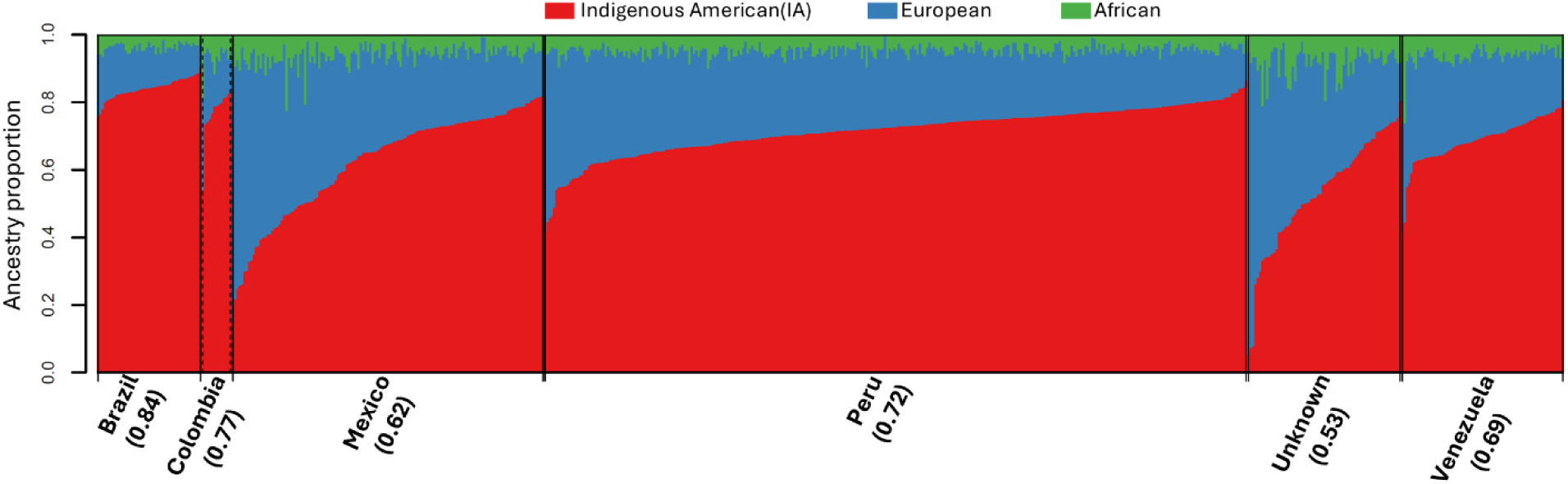
The ancestry proportions of the Latin Americans in the PAGE data by the countries of origin. The numbers below the country names represent the mean of the Indigenous American ancestry proportions. The ancestry proportions were computed with the local ancestry calls inferred by RFMix.

**Figure S10.**
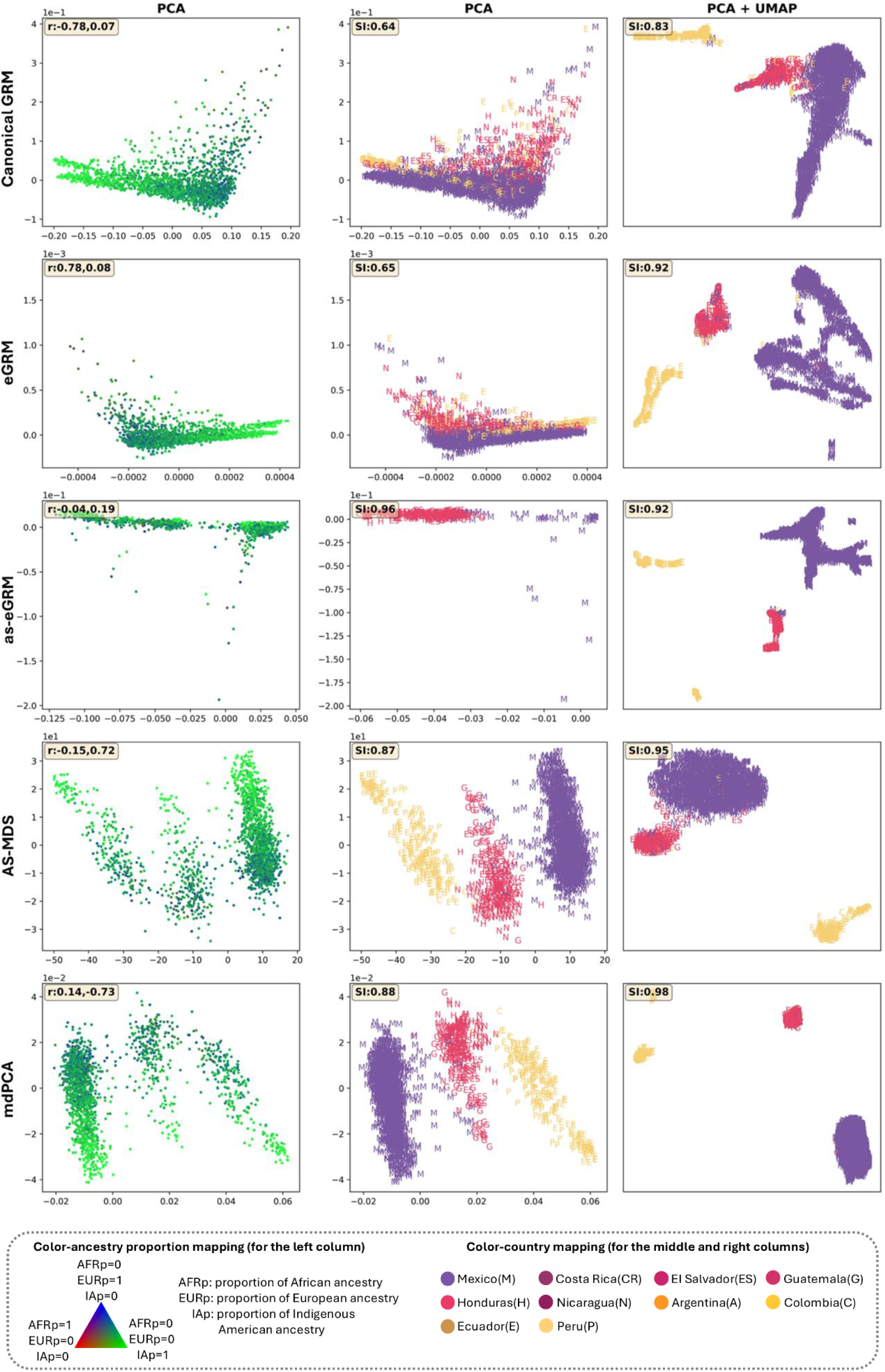
The as-eGRM replicates the Indigenous American (IA) ancestry-specific structure in the Hispanic/Latino population as demonstrated by AS-MDS using the HCHS/SOL data. PCA or PCA+UMAP result (column annotations) for each method (row annotations) when applied to a subset of 1867 HCHS/SOL individuals across all recruitment centers with estimated IA ancestry proportion > 0.5. Points represent individuals, colored by ancestry proportions (left column) or country of origin (middle and right columns, see bottom box). The r in the left upper corner of the left column represents the Pearson correlation coefficients between the proportions of Indigenous American ancestry and PC1 (the first number) or PC2 (the second number), respectively. The Separation index (SI) in the left upper corner is calculated assuming the self-reported country of origin as the true labels. Axes for UMAP plots are not labeled as distances are meaningless after UMAP transformation.

**Figure S11.**
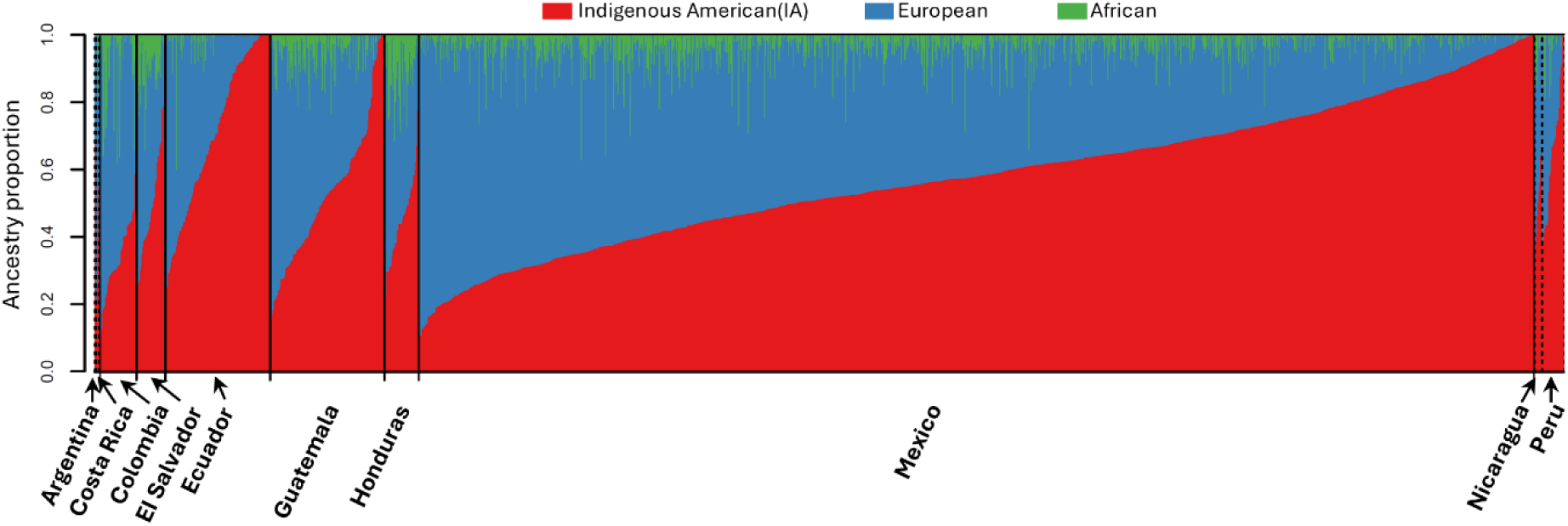
The ancestry proportions of the Hispanic/Latino individuals of Chicago recruitment site in the HCHS/SOL data. The lowest row annotates the countries where the four grandparents were self-reported to be from. The ancestry proportions were computed with the local ancestry calls inferred by RFMix.

**Figure S12.**
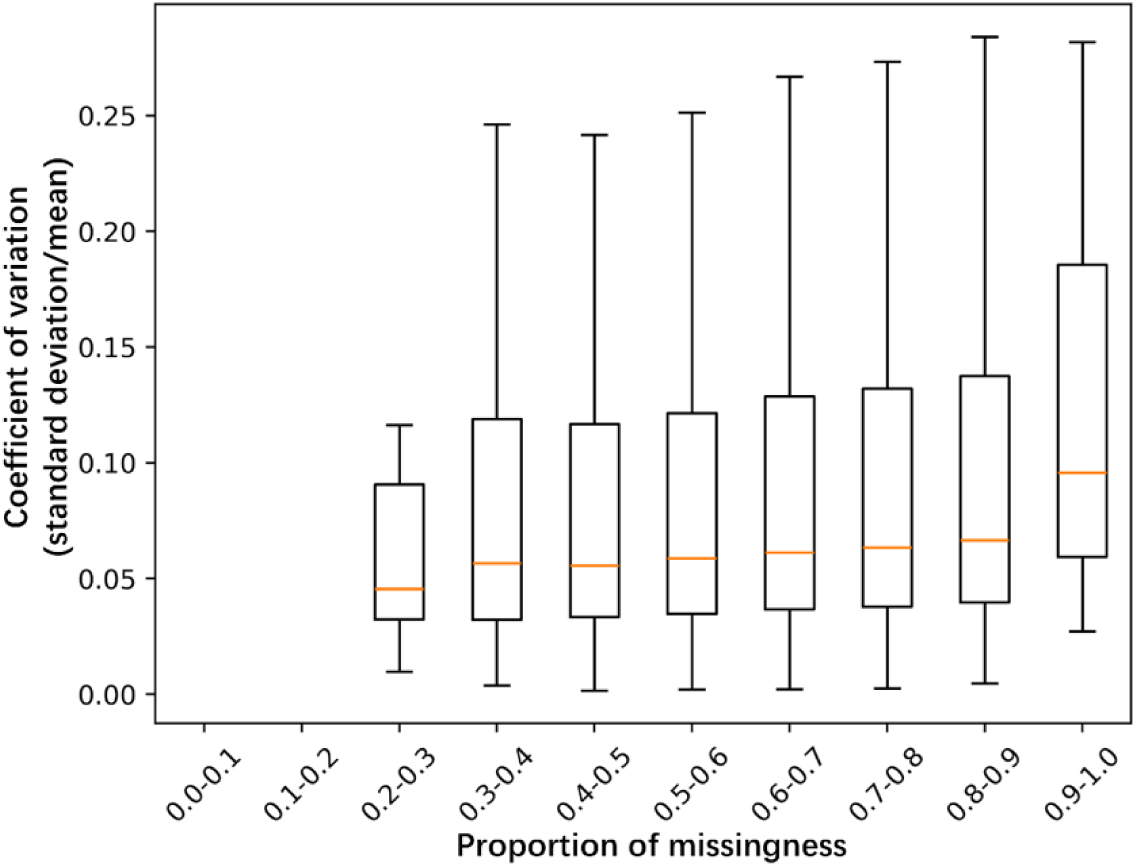
The Coefficient of variation of the relatedness estimated by as-eGRM. In data simulated by demographic history from the two-population split two-way admixture model (**Figure S2)**, we estimated the variation in the relatedness estimates by as-eGRM through 100 bootstrap samples. The distribution of the coefficient of variation as function of missingness between all pairs of individuals are shown. In general, the standard error is within 10% of the relatedness estimates themselves and empirically we have shown that as-eGRM is robust to missingness when applied to datasets with individuals across entire spectrum of ancestry proportions (Figure 5).

## Notes

### Competing Interest Statement

The authors have declared no competing interest.

